# Distinct natural syllable-selective neuronal ensembles in the primary auditory cortex of awake marmosets

**DOI:** 10.1101/2020.02.16.951194

**Authors:** Huan-huan Zeng, Jun-feng Huang, Jun-ru Li, Zhiming Shen, Neng Gong, Yun-qing Wen, Liping Wang, Mu-ming Poo

## Abstract

Marmosets are highly social non-human primates living in families. They exhibit rich vocalization, but neural basis underlying complex vocal communication is largely unknown. Here we report the existence of specific neuronal ensembles in marmoset A1 that respond selectively to distinct monosyllable or disyllable calls made by conspecific marmosets. These neurons were spatially dispersed within A1 but distinct from those responsive to pure tones. Syllable-selective responses were markedly diminished when individual domains of the syllable were deleted or the domain sequence was altered, indicating the importance of global rather than local spectral-temporal properties of the sound. Disyllable-selective responses also disappeared when the sequence of the two monosyllable components was reversed or their interval was extended beyond 1 second. Light anesthesia largely abolished disyllable-selective responses. Our findings demonstrate extensive inhibitory and facilitory interactions among syllable-evoked responses, and provide the basis for further study of circuit mechanisms underlying vocal communication in awake non-human primates.

---

Marmosets are considered to be an excellent animal model for studying neural substrates underlying complex vocal communication^1,2^. Previous brain imaging and electrophysiological studies of primate auditory systems have shown that neurons in the rostral temporal lobe show high preference for complex vocal sounds^3–5^, whereas neurons in more caudal areas such as the primary auditory cortex (A1) are well-known for their tonotopic properties, with clustered distribution into regions with preference for specific frequencies^6,7^. In addition to their frequency preference, A1 neurons are also found to be sensitive to specific spectral-temporal features of the sound, e.g. harmonicity^8^, frequency and temporal modulation^9,9^.

Electrophysiological studies in A1 of anesthetized marmosets have detected neurons that responded selectively to monosyllable Twitter^11,10^. However, it is unclear whether A1 neurons could selectively respond to all natural calls, including both mono- and disyllables, and whether syllable-evoked responses was only due to neurons’ sensitivity towards specific local spectral-temporal feature of the sound or requiring global temporal organization of various sound components, such as the sequence and interval of monosyllables within the disyllable. It is thus important to perform simultaneous recording of the activity from large A1 neuronal populations in the same marmoset. Such recording needs to be conducted in the awake state, since anesthesia is known to greatly reduce neuronal activity in the cortex.

In this study, we have achieved two-photon fluorescence imaging of large-populations of A1 neurons in un-anesthetized marmosets by acute loading of Ca^2+^-sensitive fluorescent dye Cal-520AM. This method allows rapid labeling of a much larger proportion of neurons than that currently could be achieved by genetic expression of GCaMP6. Using this method, we have identified within conventional tonotopic regions of A1 subtantial populations of neurons that respond selectively to different conspecific mono- and disyllables but not to pure tones. Further studies focusing on disyllable-selective neurons showed that their responses are sensitive to the sequence and interval of monosyllable components, characters of vocal sound processing. These disyllable-selective responses were found only for naturally occurring but not artificially constructed disyllables, and were completely abolished by light anesthesia. These findings established the existence of substantial syllable-selective neuronal ensembles in A1 of awake marmoset, pointing to complex vocal sound processing in the early stage of the auditory system.

## Results

### Two-photon Ca^2+^ imaging of neuronal activity in A1

We simultaneously monitored the activity of a large population of A1 neurons in head-fixed awake common marmosets by fluorescence Ca^2+^ imaging. The A1 area was first identified based on its tonotopic organization, as revealed by imaging intrinsic optical signals in anesthetized marmosets (Extended Data Fig. 1)^6,7^. Synthetic Ca^2+^-sensitive dye Cal-520AM (ref. 6) was then loaded into a specific subregion of A1 (2-8 kHz areas, areas >8 kHz were buried in the sulcus) to label layer 2/3 neurons (see Methods). Two-photon Ca^2+^ imaging of neuronal activity in response to various natural syllables (Extended Data Fig. 2a, b; Supplementary Video 1) was performed 2 hours after dye loading when the marmoset regained wakefulness, and the recording normally lasted for 3 hours.

In an alternative approach, we injected tetracycline (Tet)-activated AAV vector expressing genetically encoded Ca^2+^-indicator GCaMP6f (ref. ^11^) into A1 and performed imaging more than 4 weeks after injection and 3 days after Tet feeding (Extended Data Fig. 2c, d; see Methods; Supplementary Video 2). Although GCaMP6f expression was detectable in a lower proportion of neurons as compared to Cal-520AM, this approach allowed repetitive recording from the same neuronal populations, showing the stability of syllable-evoked neuronal responses in the same marmoset over a duration of about one week (Extended Data Fig. 2e, f). Both imaging approaches yielded similar results, and the data were pooled in some analyses.

### Consepecifc syllable-selective responses

To detect neurons that could respond selectively to the same call syllables made by conspecific marmosets, we performed two-photon imaging of neuronal Ca^2+^ signals in A1 sub-regions of two marmosets (M_a_ and M_b_) that were acutely loaded with Cal-520AM, and monitored population neuronal responses to Phee (P), Twitter (Tw) and TrillPhee (TrP) recorded from three other marmosets (M_1_, M_2_, M_3_; 3 calls for each syllable, 27 calls in total, spectrograms shown in Extended Data Fig. 3a, Supplementary Video 3-5). Analysis of the spectral-temporal properties of the 27 call samples by principal component analysis and bandwidth Wiener entropy showed distinct clustered distribution of calls of the same syllable category, despite the fact that there were substantial differences in the duration and spectral-temporal properties among calls made by different marmosets (Extended Data Fig. 3b).

As illustrated in Fig. 1a, each of the 3 example neurons from M_a_ showed selective responses to the same syllable made by two or three different marmosets. We defined neuronal responses to be syllable-selective when the mean Ca^2+^ fluorescence change (ΔF/F) evoked by a syllable category (n = 9, 3 calls from each marmoset) was significantly higher (at a level larger than 5-fold) than those evoked by the two other syllable categories (*P* < 0.05, ANOVA; see Methods). Average responses (ΔF/F) of all syllable-selective neuronal ensemble in marmoset M_a_ evoked by 27 syllable samples were depicted by the heat map in Fig. 1b. Summary of all data from M_a_ and M_b_ showed consistent selectivity of the same neuronal ensemble towards conspecific syllables (Fig. 1c).

**Fig. 1.**
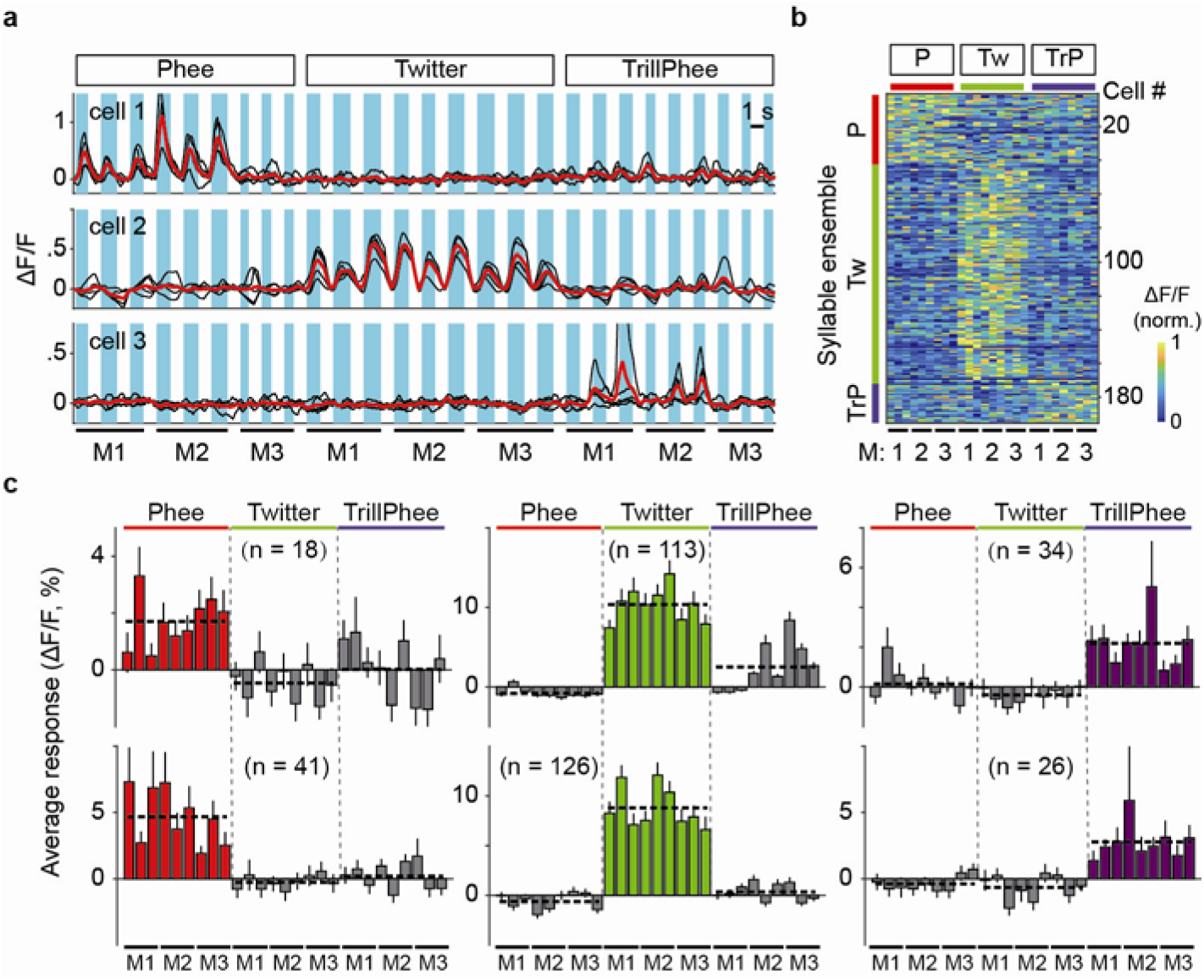
A1 neurons in awake marmosets selectively responded to conspecific syllables. **a**, Fluorescence changes (ΔF/F) in 3 example cells (in marmoset M_a_) evoked by 27 conspecific test syllables. Note that each cell responded selectively to either Phee, Twitter or TrillPhee calls (n = 3) from 3 different marmosets (M_1_, M_2_ and M_3_). Black traces: single trials (n = 5); red traces: average; shading: syllable duration. **b**, Heat map for all syllable-selective neurons in one marmoset (M_a_) that was exposed to three syllable categories as in **a**. Each horizontal line depicts average amplitude of ΔF/F (from 5 trials), with 3 representative calls from each marmoset for each syllable. The cells were sorted into three syllable ensembles, based on the syllable they exhibited the highest mean ΔF/F amplitude. The amplitude is coded in color by the scale shown on the right. **c**, Average response amplitudes of neuronal ensembles that selectively responded to Phee, Twitter and TrillPhee (error bar, SEM; n= total number of neurons examined). Data were from marmoset M_a_ (bottom) and M_b_ (top) respectively. Dash horizontal lines: mean response of each ensemble.

We have examined whether neurons responding selectively to syllables could also respond to other non-syllable variables, including call duration, bandwidth, Wiener entropy, amplitude modulation and caller identity (M_1_, M_2,_ or M_3_). Generalized linear model was used to test syllable-selective neuronal ensembles identified in the experiment described in Fig. 1. We indeed found many of these neurons were significantly modulated by one or more of non-syllable variables (Extended Data Fig. 4, see details in Methods). However, a substantial number of them (27/193, 14.0%, M_a_; 22/165, 13.3%, M_b_) showed an exclusive selectivity to syllables and not to other variables, indicating the existence of neuron ensembles in A1 that are purely syllable-selective, without being affected by other acoustic factors and caller identity (Extended Data Fig. 4a, M_a_; Extended Data Fig. 4f, M_b_). Furthermore, using multidimensional scaling to visualize neuronal representations of syllables and non-syllable variables (Extended Data Fig. 4b-e, M_a_; Extended Data Fig. 4g-j, M_b_), the exclusive syllable-selective neurons showed three distinct clusters. However, such clustering was not present when neurons selective to non-syllable variables were examined, with Phee and TrillPhee neuron clusters undistinguishable in both marmosets. Thus, syllable-selective neurons we have observed consist of neurons that responded exclusively to distinct syllables as well as neurons whose syllable-selective responses were significantly modulated by acoustic and caller identity variables.

### A1 neuronal responses to four standard syllables

To further investigate the population characteristics and spatial distribution of syllable ensembles in A1, we adopted four most common syllables for the standard test (three monosyllables: Phee, Twitter, Trill, and one disyllable TrillPhee; Fig. 2a). Many A1 neurons responded selectively to a specific syllable (examples in Fig. 2b), and all neurons showing syllable selectivity were sorted according to the time of the peak ΔF/F signal to obtain the activity profile map, revealing clear syllable-selective neuron ensembles within the imaged A1 area of marmoset M_c_ (Fig. 2c). Data for two other marmosets M_a_ and M_d_ are shown in Extended Data Fig. 5a. Notably, within each ensemble, the peak response time of neurons tiled the entire syllable duration (from 100s ms to >1 s), with more neurons reached peak firing near the end of the syllable sound (Fig. 2c). As discussed later, this temporal tiling of neuronal responses over the range of ~ 1 s is critical for interval timing in facilitary and inhibitory interactions among syllable-evoked responses. The relative sizes of syllable ensembles appeared to be different among the three marmosets examined. Notably, Phee neurons did not respond to TrillPhee that contains the Phee component, indicating that monosyllable-evoked responses are sensitive to the temporal context of the syllable.

**Fig. 2.**
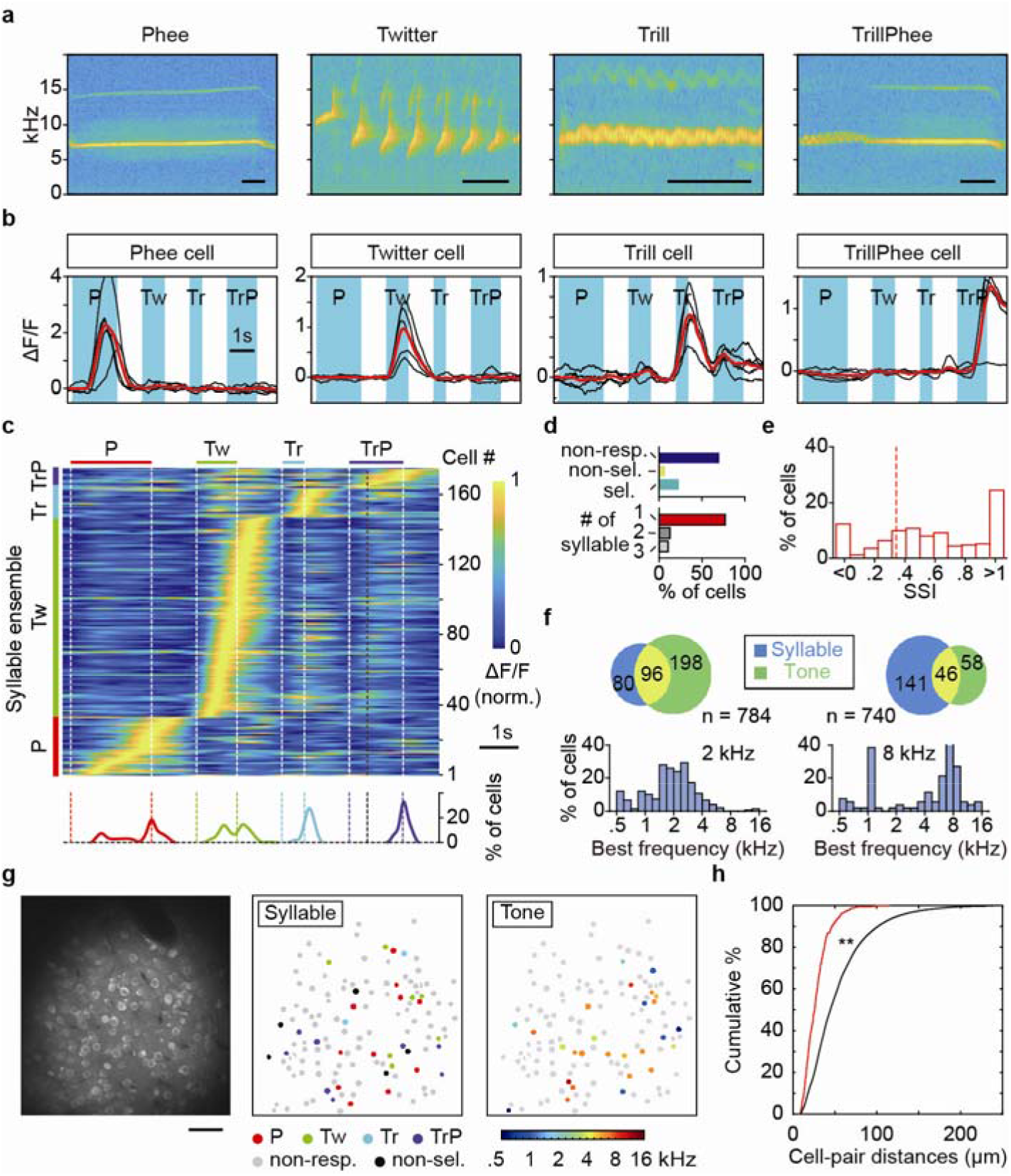
Analysis of syllable-selective cells in awake marmoset A1. **a**, Representative spectrograms of 4 standard test syllables. Bars: 0.2 s. **b**, Fluorescence changes (ΔF/F) in 4 syllable-selective cells in A1, recorded from marmoset M_c_ that was loaded with Cal-520AM. Black traces: single trials (n = 5); red traces: average; cyan shading: syllable duration. **c**, Heat map for the activity of all syllable-selective cells in M_c_, with the cell sorted in an order based on the time of peak ΔF/F. White dashed lines: syllable onset and offset; black dashed line: boundary of Trill and Phee components of TrillPhee. Bottom: traces depicting percentages of cells that had different peak-response times within each syllable ensemble. **d**, Statistics on syllable-selective cells recorded from 24 imaging fields in 3 marmosets (M_a_, M_c_ and M_d_) labeled with Cal-520AM. Top: among all cells recorded (n= 2,891), the percentages of cells that were unresponsive, responsive but not syllable-selective and syllable-selective. Bottom: the percentages of cells showing syllable selectivity to 1, 2, or 3 syllables. **e**, Syllable Selective Index (SSI) of all syllable-selective cells. Red dashed line: SSI = 0.33 (2-fold preference). **f**, Top: Venn chart of the number of syllable-selective neurons and pure-tone responsive neurons, with the overlap representing the number of cells with both types of responses. Bottom: the percentage of pure-tone responsive neurons showing different best frequencies (left, 2-kHz area; right, 8-kHz area). **g**, Left, an image of Cal-520AM fluorescence at a recorded region. Bar: 50 μm. Middle: spatial distribution of all cells in the imaging field, with cell response properties coded in colors. Right: tonotopic properties of the imaging field. **h**, Cumulative percentage plot of nearest-neighbor distances for cells of the same syllable selectivity (red line), and for all cells regardless of syllable selectivity, obtained by bootstrap analysis (black line, see Methods). The difference between two distributions is significant at *P* < 0.001, *Kolmogorov-Smirnov test*).

Among all A1 neurons examined in three marmosets (M_a_, M_c_ and M_d_) loaded with Cal-520AM, we found that ~23% (674/2,891) showed significantly higher mean response amplitude to one or more syllables (*P* < 0.05, ANOVA). A small fraction of them (75/674) exhibited similar mean response amplitudes for 2 or 3 syllables (*P* > 0.05, t-test; Fig. 2d, Extended Data Fig. 5b), and a few showed positive ΔF/F to one syllable but negative ΔF/F to another (Extended Data Fig. 5c). Among syllable-selective neurons, Twitter neurons were most common, followed by Phee, TrillPhee and Trill neurons (Extended Data Fig. 5b). Twitter neurons were also the prominent type of syllable-responsive neurons observed in electrophysiological studies of anesthetized animals^10^. Quantification by Syllable Selectivity Index (SSI, see Methods) showed that most syllable-selective neurons exhibited high selectivity (with SSI > 0.33, or 2-fold difference, Fig. 2e).

### Neuronal populations selectively responding to pure tones or syllables

The A1 sub-regions chosen for the above experiments had tonotopic preference for either ~2 or ~8 kHz, as determined by imaging intrinsic optical signals. Neurons were considered pure tone-selective based on conventional criteria^6^, and all A1 neurons within the imaging field were examined for the responses evoked by pure tones ranging from 0.5 to 16 kHz (5 frequency samples per octaves). Syllable-selective neurons were determined by the criteria described above. Our measurements of all Cal-520AM-labeled neurons (n = 784 in M_a_ and 740 in M_c_) showed that only a small percentage of neurons responded selectively to both syllables and pure tones (Fig. 2f; 8-kHz area, 6%; 2-kHz area, 12%). Furthermore, the percentage of syllable neurons was higher than pure-tone neurons in the 8-kHz area (25% *vs*. 14%), and the opposite was found for the 2-kHz area (22% *vs*. 38%, Fig. 2f). Further examination of the spatial distribution of different syllable- and pure tone-selective neurons within the same imaging fields showed that neurons belonged to different syllable ensembles appeared to be spatially intermingled (Fig. 2g, Extended Data Fig. 6). However, the nearest-neighbor distances for neurons of the same ensemble were on the average smaller than those for neurons randomly sampled from different ensembles (Fig. 2h, *P* < 0.001, bootstrap analysis), suggesting some spatial clustering of neurons of the same syllable ensemble.

The syllable-selective neurons were found to be relatively sparse and dispersed within A1 tonotopic areas, unlike the clustering of face-selective neurons in the inferior temporal cortex. The tendency of closer apposition among neurons within the same syllable ensemble may reflect intracortical circuit organization underlying the syllable-selective responses. While the proportion of syllable-selective neurons in A1 within each imaged field was comparatively low (e.g., <10% for Twitter cells), the estimated total population within A1 for each syllable ensemble could reach tens of thousands. Taking Twitter neuron ensemble for example, the volume and neuronal density of marmoset A1 is estimated to be 8.18 mm^3^ and 78080 neurons/ mm^3^ acording to a previou study^12^, the overall neuron number of A1 is 638,690, among which Twitter neuron number is ~63,869 (~ 10%).

### Response properties of neuronal ensembles for disyllables

Further measurements of neuronal responses to two disyllables TrillPhee (TrP) and TrillTwitter (TrTw) showed that response onset generally occurred after the appearance of the second monosyllable component (Fig. 3a, b, cell 1), and only a few of these disyllable neurons responded weakly to isolated monosyllable Phee or Twitter (Fig. 3a, b, cell 2). Thus, the immediate prior presence of Trill greatly enhanced the neuronal response toward Phee or Twitter, indicating the facilitary action of Trill on these disyllable neurons. Monosyllable-selective Phee and Twitter neurons did not respond to Phee and Twitter components within the TrillPhee and TrillTwitter disyllables (see Extended Data Fig. 5c, e), indicating the inhibitory action of immediately preceding Trill on the responses of these monosyllable neurons. Activity heat maps of all TrillPhee neurons (Fig. 3c, M_a_, M_c_, M_d_; Cal-520AM-labeled) and TrillTwitter neurons (Fig. 3d, M_a_, GCaMP6f-labeled), as well as Trill, Phee, or Twitter monosyllable-selective neurons, showed that the size of disyllable ensembles could be as large as that of monosyllable ensembles.

**Fig. 3.**
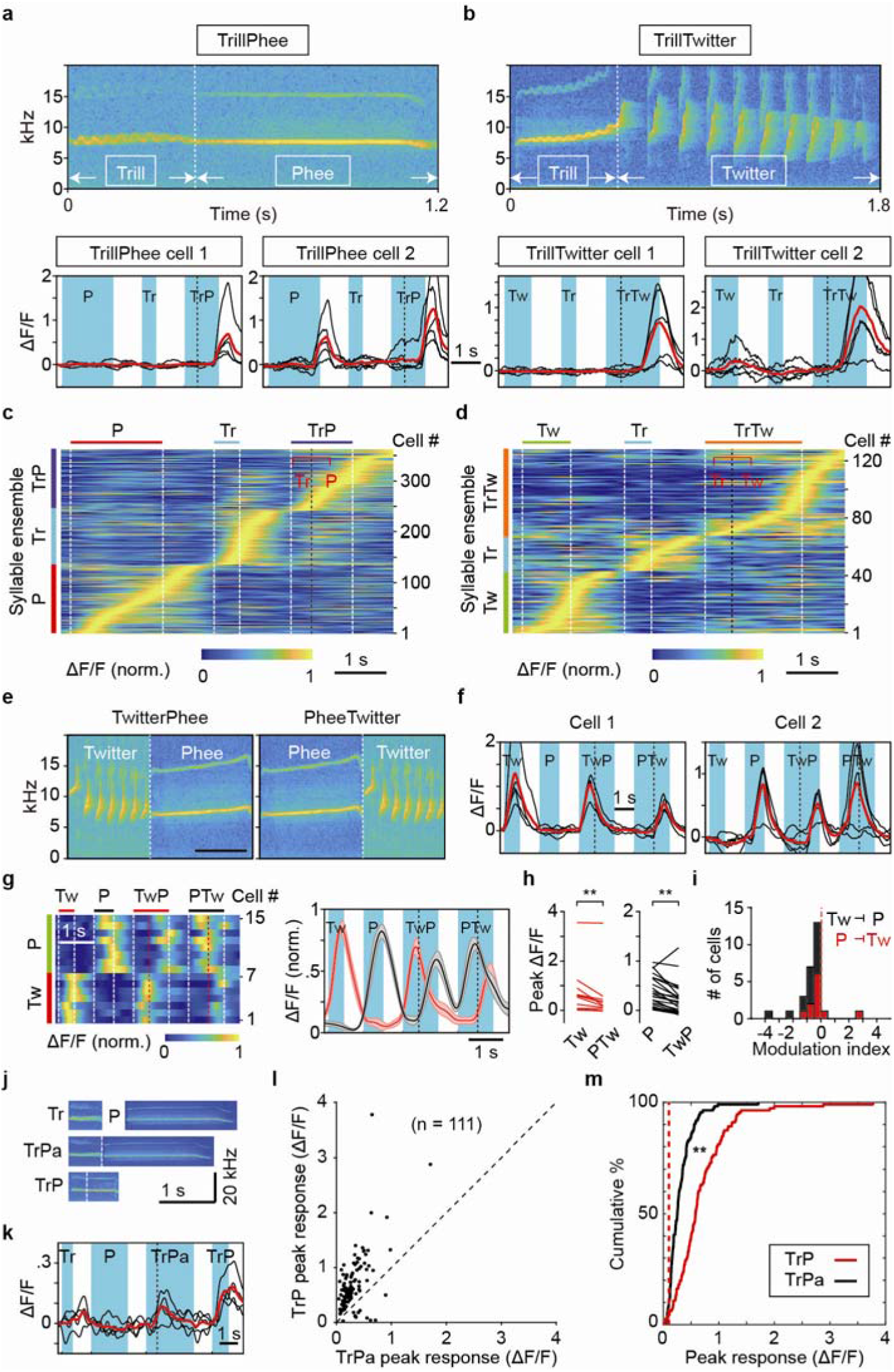
Properties of disyllable-selective cells. **a, b**, Spectrograms of disyllables TrillPhee and TrillTwitter, and selective responses of two example cells for each disyllable. **c, d**, Heat maps of the activity of all cells selectively responding to disyllables (**c**, M_a_, M_c_, M_d_, Cal-520AM-labeled; **d**, TrillTwitter, M_a_, GCaMP6f-labeled) and monosyllable components (Trill, Phee, Twitter). Black dashed line, boundary of monosyllable components. **e**, Spectrograms of novel disyllables TwitterPhee and PheeTwitter. Bars: 1 s. **f**, Single trials (black lines, n = 5) and mean (red line) evoked by Twitter, Phee, TwitterPhee and PheeTwitter in two example cells. Dashed line, boundary of Phee and Twitter. **g**, Heat map of normalized responses to monosyllables and artificial disyllables (left) for example cells that show selective response to Twitter (n = 7) and Phee (n = 8), and their responses to artificial disyllables. Right: normalized mean ΔF/F (± SEM) induced by monosyllables and artificial disyllables for all cells of the Twitter (red) and Phee (black) ensembles, corresponding to the heat map on the left. Note that both Twitter and Phee cells responded to artificial TwitterPhee and PheeTwitter with reduced amplitudes. **h**, Comparison of the peak ΔF/F values for individual neurons within the Twitter (n=11) and Phee (n = 26) ensemble, between responses to isolated monosyllables and those to the same monosyllable within artificial disyllables (**, *P* < 0.01, paired t test). **i**, The inhibitory effect of one monosyllable to another that followed immediately, as quantified by the modulation index (MI) that represents fractional changes in the peak ΔF/F of monosyllable-evoked responses (see Methods). Note that MIs were predominantly negative for both Twitter and Phee neurons. **j**, Spectrograms of natural Trill, Phee and artificial TrillPhee (TrPa) made from natural Trill and Phee, and a natural TrillPhee. All syllables are from the same marmoset M_2_. **k**, Single trials (black lines, n = 5) and mean (red line) evoked by Trill, Phee, TrPa and natural TrillPhee in an example cell. Dashed line, boundary of Trill and Phee. **l**, Responses to natural TrillPhee and TrPa of 111 neurons. **m**, Cumulative percentages of neurons that responded to natural and artificial TrillPhee with diffenrent amplitudes. Red dashed line, value of 0.1 in ΔF/F. The difference between two distributions is significant at *P* < 0.001, *Kolmogorov-Smirnov test*).

We have also constructed artificial disyllables by linking two natural monosyllables Twitter and Phee from the same marmoset M_0_ (Fig. 3e). These novel disyllables TwitterPhee and PheeTwitter were never recorded in our marmoset colony. We found that these novel disyllables failed to elicit any disyllable-selective response. All neuronal responses appeared to be evoked by the monosyllable Twitter or Phee (Fig. 3f, M_a_ and M_d_), and the peak amplitudes of these artificial disyllable-evoked responses were slightly lower than those evoked by isolated Phee or Twitter (Fig. 3g-i), again implicating the inhibitory action between monosyllables. We also constructed artificial TrillPhee by joining two randomly sampled monosyllable Trill and Phee recorded from M_2_, and found that such artificial TrillPhee could also evoked significant selective responses in many A1 neurons, but the response amplitudes were consistently lower than those evoked by the natural TrillPhee made by the same animal, as shown by the example neuron (Fig. 3k) and all neurons examined (n =111, Fig. 3l,m).

Taken together, these results on artificial disyllables suggest that disyllable-selective neurons in A1 are designed for detecting natural disyllable calls, through natural selection or auditory experience, or both.

### Effects of disyllable domain deletion, sequence alteration, and interval extension

Are syllable-selective responses of A1 neurons due to unique spectral-temporal property of a specific sound domain within the syllable? We address this question by focusing on the disyllable TrillPhee which has a more complex spectrogram. In two GCaMP6f-expressing marmosets, we first performed “domain deletion” experiments, in which three separate domains of the disyllable TrillPhee (D_1_: Trill; D_2_: Trill-Phee junction; D_3_: Phee) were sequentially deleted (Fig. 4a). We found that deleting either one or two domains of the disyllable markedly reduced the response of disyllable neurons (Fig. 4b, 2 example cells; Fig. 4c, all 10 cells recorded in M_a_ and M_b_). This indicates that TrillPhee responses were due to global rather than local spectral-temporal property of the disyllable. In further “domain sequence alteration” experiments, whereby the three TrillPhee domains were all present but their temporal sequence altered in 5 different ways, we found that any alteration of the natural sequence (D_1_/D_2_/D_3_) resulted in marked reduction of evoked responses (Fig. 4e, f). Thus, both the presence of all domains and their proper temporal sequence are critical, implicating sequence-specific integration of information on different sound components by the responsive neurons. The importance of temporal sequence of sound components was further confirmed by the finding that reversing the Trill/Phee sequence into Phee/Trill completely abolished the TrillPhee-selective responses in all 9 of TrillPhee neurons examined (Fig. 4g-i, marmoset M_a_).

**Fig. 4.**
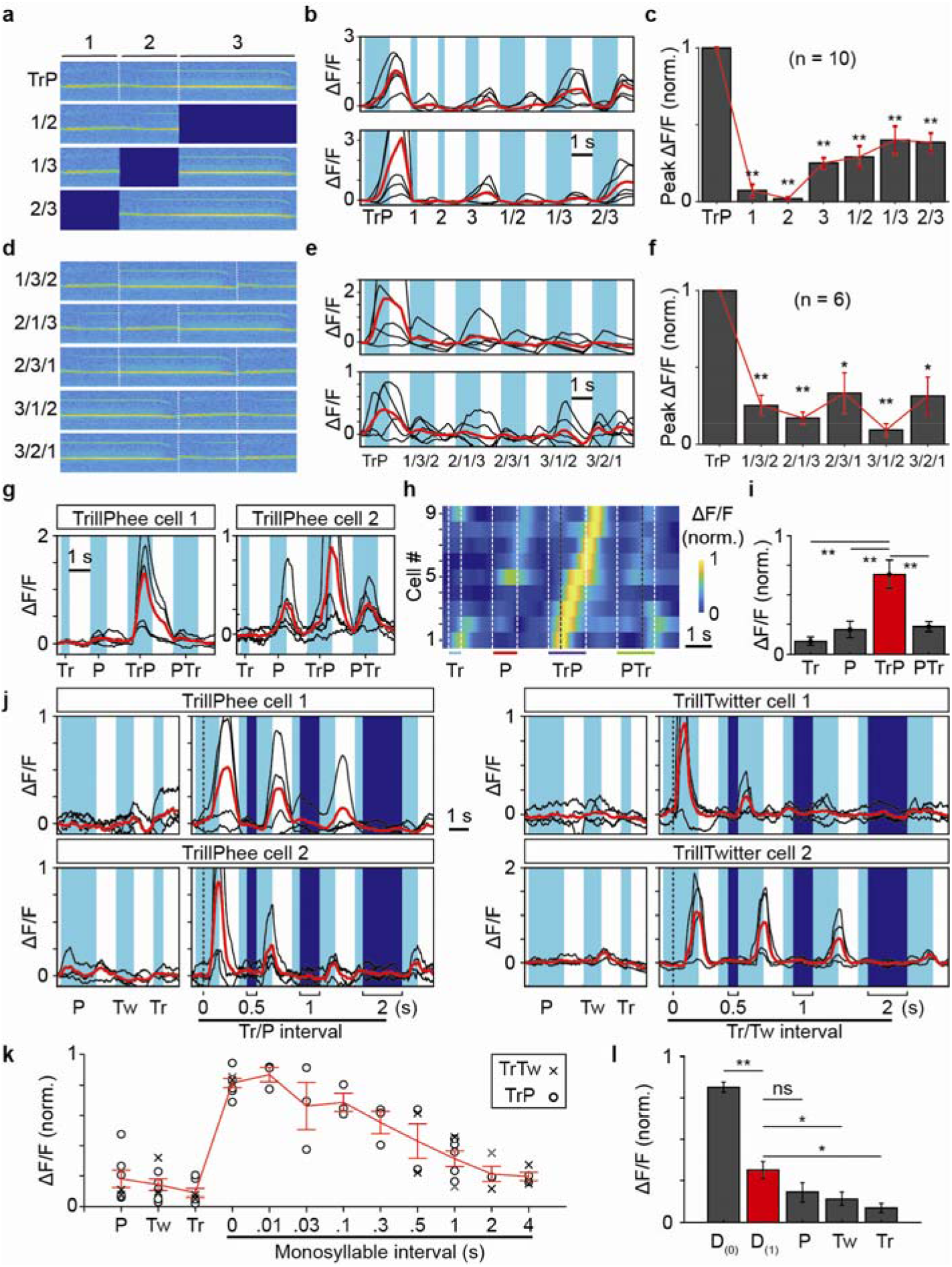
Experiments on “domain deletion”, “domain sequence alteration”, and monosyllable sequence reversal and interval extension. **a**, Spectrograms of a complete TrillPhee and domain-deleted TrillPhee, in which one of three domains (1, Trill; 2, Trill/Phee junction; 3, Phee) was deleted. **b**, Two example TrillPhee neurons responding to the complete TrillPhee and one or two TrillPhee domains. **c**, Summary of normalized peak ΔF/F values for all 10 TrillPhee cells examined in domain-deletion experiments. **d**, Spectrograms of TrillPhee with domain sequence alteration, based on three domains defined in **a. e**, Two examples of TrillPhee cells responding to complete TrillPhee and 5 different domain sequence-altered TrillPhee. **f**, Summary of normalized peak ΔF/F values for all 6 TrillPhee neurons examined in domain sequence alteration experiments. **g**, Two examples of TrillPhee neurons showed complete loss of disyllable selectivity when the Trill/Phee sequence was changed to Phee/Trill. **h**, Heat map of 9 TrillPhee neurons examined in the “reverse sequence” experiment, showing responses to TrillPhee but not PheeTrill. The ΔF/F value was normalized for each cell. **i**, Summary of average peak ΔF/F values for all cells shown in h (**, *P* < 0.001, paired t test). **j**, Two example cells with disyllable-selective responses to natural disyllables (Left, TrillPhee; Right, TrillTwitter) and reconstructed disyllables with an interval of 0.5, 1, or 2 s between two component monosyllables, together with their responses to isolated monosyllables Trill, Phee, and Twitter. **k**, Summary of all data on responses evoked by reconstructed disyllables with extended intervals from 0.01 to 4 sec (n = 3-7 cells each) and by 3 isolated constituent monosyllables, recorded from marmoset M_a_ expressing GCaMP6f. Red curve: averages at all intervals, with data points depicting normalized peak value of ΔF/F for two disyllables. **l**, Averages of normalized peak ΔF/F values for data in **k**, for natural disyllable [D_(0)_], extended disyllable with 1-s interval [D_(1)_], and three constituent monosyllables (n = 7 cells; paired t test; **, *P* < 0.001; *, *P* < 0.01; ns, *P* > 0.05).

In addition to domain sequence specificity, we further examined reconstructed disyllables in which the interval between monosyllable components was extended from 10 msec up to 4 sec. The responses were found to decline around an interval of ~100 ms and largely disappeared beyond 1 sec (examples, Fig. 4j; summary, Fig. 4k and 4l). Disyllables with over-extended intervals between monosyllable components still triggered weak responses in some disyllable neurons (Fig. 4j), with amplitudes similar to that evoked by isolated monosyllables. Thus, normal disyllable-selective responses require not only proper sequence of the monosyllable components, but also their temporal proximity within ~1 sec.

In a parallel experiment, we monitored the activity of Phee-selective monosyllable neurons with the imposition of a preceding Trill (isolated from a TrillPhee call) at an interval of 0, 0.5, 1, or 2 sec, and found that the suppression effect of preceding Trill gradually reduced as the interval of Trill/Phee was increased (Extended Data Fig. 7). Thus, monosyllable-induced suppression is also interval-dependent.

### Effects of anaesthesia on syllable-selective responses

Many previous studies of auditory processing in non-human primates were performed in anesthetized preparations^13–15^. In this study, we adopted a fentanyl cocktail for light anesthesia^16^, under which pure-tone responses were still robustly evoked in A1^6^, and the overall level of Cal-520 AM fluorescence remained largely unchanged (Fig 5a). We found that this anesthesia modulated the responses of both monosyllable and disyllable neurons. Many monosyllable neurons still exhibited syllable-selective responses with lower amplitudes, but their temporal profiles were altered (Fig. 5b and 5c). Notably, a large proportion of TrillPhee neurons became completely non-responsive to TrillPhee (Fig. 5b and 5c). Comparison of response profiles of the same ensemble of neurons before and during anesthesia showed that the anesthesia reduced the amplitude, duration, and syllable selectivity of the responses (Fig. 5d-f).

**Fig. 5.**
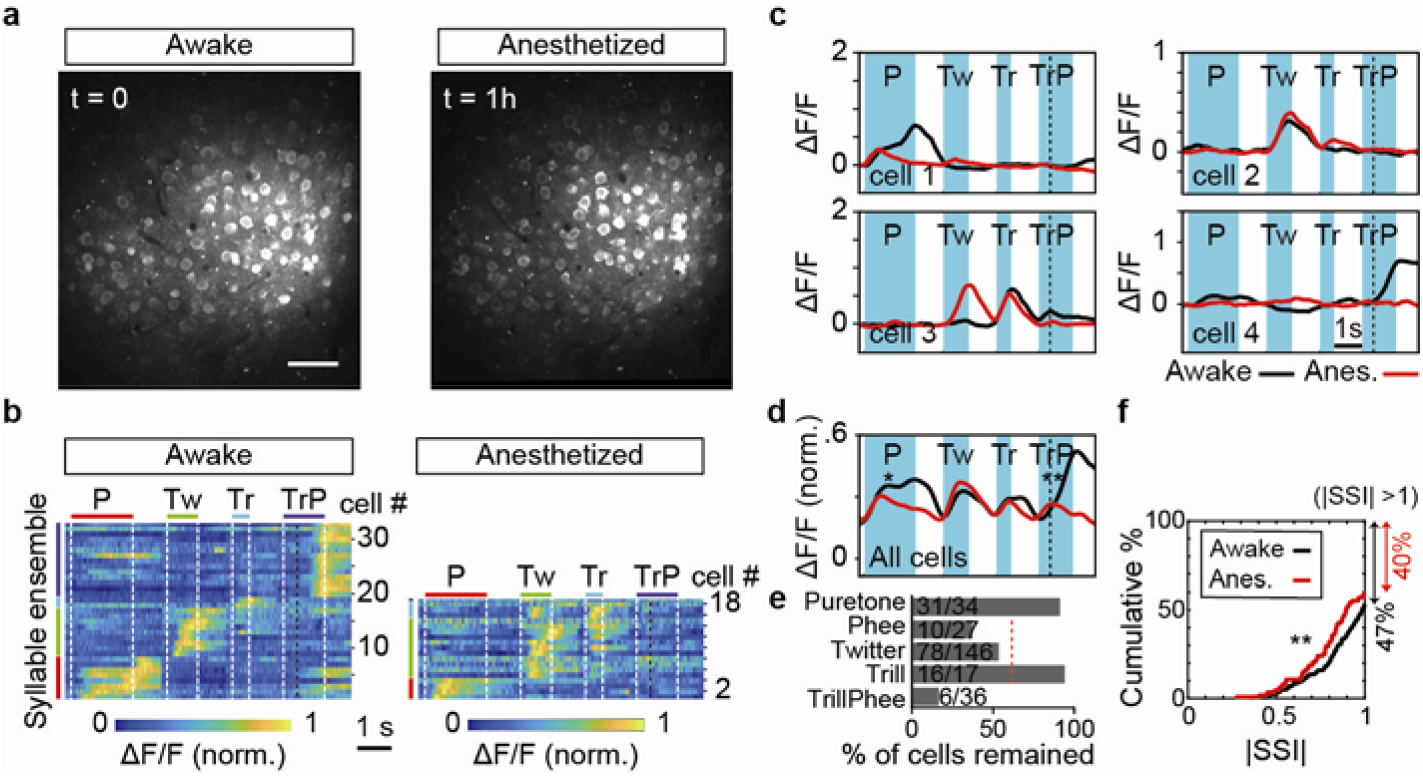
Anesthesia reduced syllable-selectivity. **a**, Images of Cal-520AM fluorescence (averaged over 2 min) at a recorded region in marmoset M_d_, before (left) and 1-hour after (right) induction of light anesthesia with a fentanyl cocktail. Bar, 50 μm. **b**, Heat maps of the activity of syllable-selective cells within an example imaging field (shown in **a**) in awake state and 1-hour after anesthesia. Note that TrillPhee neurons largely disappeared after anesthesia (only two remained). **c**, Four example cells depicting syllable-selective responses shown in **b**, with each trace depicting averaged signals from 5 trials. **d**, Summary of all data on syllable-selective cells (n = 62, 3 imaging fields, M_d_) before (black) and 1-hour after (red) anesthesia, shown by the average traces of ΔF/F. The mean ΔF/F values after anesthesia was determined based on the normalization used for the same neuron in the awake state. Significant differences were found for disyllable TrillPhee and monosyllable Phee neurons (P, *P* < 0.01; TrP, *P* < 0.001; Tw, Tr, *P* > 0.05; t test). **e**, The percentage of total cells remained showing pure-tone, Phee, Twitter, Trill and TrillPhee responses, 1-hour after anesthesia induction. Red dashed line, mean value of monosyllable Phee, Twitter, and Trill. Data were from M_b_ and M_d_. **f**, Cumulative percentage plot of the distribution of absolute Syllable Selective Index (|SSI|) values for all syllable-selective cells before (black line, n = 226, 47 % of neurons showing |SSI| > 1) and 1-hour after (red line, n = 112, 40 % of neurons showing |SSI| > 1) anesthesia induction. Data were from M_b_ and M_d_. The difference between two distributions (include all cells) is significant at *P* < 0.05, *Kolmogorov-Smirnov test*).

## Discussion

Auditory processing in A1 is characterized by the tonotopic organization and spectral-temporal selectivity of neuronal responses^17,18^, presumably involving feed-forward thalamocortical inputs and intracortical processing by local circuits^19–21^. Here we show that, in marmoset A1 tonotopic regions comprising neurons predominently tuned to specific sound frequencies, there are substantial populations of neurons specifically devoted to syllable processing. To determine whether neurons are selective to natural syllables or disyllables, we first examined the responses of each A1 neuron to the same syllable calls from different animals. We found that, despite some dispersion of spectra-temporal properties of the same syllable call made by different marmosets, the same selective response pattern was evoked in the same neuron, indicating invarience of the responses to conspecific syllables. Further studies using standard syllables provided evidence for complex suppressive and facilitory processing in syllable-evoked responses. First, we found that responses of monosyllable-selective neurons were suppressed by the presence of other preceding monosyllables, indicating suppressive interactions among monosyllables. Second, the requirement of the specific sequence of syllable domains and the restricted interval between monosyllable components within the disyllable implicates well-orchestrated facilitory modulation. Finally, the high susceptibility of disyllable-evoked responses to disruption by light anesthesia is consistent with the presence of polysynaptic signaling and top-down regulation, which are known to be more vulnerable to anesthesia^22,23^.

An important issue of vocal communication is the processing of temporal sequence and time interval of sound units that could span timescales from milliseconds to seconds^24^. Previous studies have shown that selectivity for conspecific sounds is present in the avian primary auditory forebrain, and spectral-temporal features of the sound could account for the neuronal responses. In marmoset A1, temporal compression, extension or reversion of marmoset Twitter calls greatly diminished neural firing evoked by natural Twitter^10^, indicating the importance of temporal feature of the sound. Our findings are in line with these previous reports and further showed that not only the spectral-temporal structure of the sound over hundreds of millisecond timescale is important, the sequence of sound components over a temporal window of seconds is also critical. Notably, syllable sound processing requires a continuous coding of sound information by responsive neurons. Such prolonged coding could be achieved by syllable-specific neuronal ensemble with peak response profiles that tiled over a period of seconds as shown by our population recording data, allowing continuous coding of the global spectra-temporal property as well as the sequence of sound components of the syllable.

A major finding of this study is the temporal context within which the syllable sound occurs, as shown by the suppressive and facilatory actions among temporally conjuctive monosyllables. For example, the immediate prior presence of Trill suppressed Phee-evoked responses of Phee-selective neurons. Such suppression disappeared when the interval between Trill and Phee was extended beyond 1 s. Such suppression could be mediated by Trill neuron-activated interneurons that provide extended inhibitory inputs to the Phee-selective neurons over a period up to 1 s, covering the entire duration of the Phee sound via temporal tiling of Trill neuron responses. On the other hand, we found that in disyllable-selective TrillPhee neurons, the immediate prior presence of Trill appeared to provide facilitory action on the neuron’s response to the subsequent Phee component. This could be accomplished by the Trill-neuron ensemble activation that causes disynaptic disinhibition of Phee evoked responses, under the condition that TrillPhee neurons are under strong inhibition that prevents their response to Phee in the absent of preceding Trill. It remains to be further determined whether these actions involve neural circuits in A1 or other regions of the auditory pathway, or both. A study using functional magnetic resonance imaging with macaques has shown preferential activity in anterior auditory fields for species-specific vocalization and vocal identify of conspecific individuals^25^. Neuronal representation of specific syllables in A1 could serve as building blocks for further circuit computations of more selective representations in higher cortical regions.

Vocal communication has been extensively studied using songbirds^26–28^, rodents^29,30^, and non-human primates^2,31^. In birds, neurons in tonotopic organized primary auditory field and caudal hyperstriatum ventral region (cHV) show extremely selective responses to the bird’s own song but not conspecific songs by others^32^. In mouse, neurons in the inferior colliculus and some auditory cortical regions respond robustly to ultrasonic vocalization^29^. Studies in humans and non-human primates have shown that neurons sensitive to conspecific vocal sounds exist in many regions of superior temporal plane^31^, including A1. Thus, syllable-selective responses in A1 could reflect activity of down- or upstream regions of the auditory pathway. Alternatively, these A1 syllable-selective neurons could be the main site of information processing underlying syllable recognition. Further experiments examining the effect of silencing activity in different brain regions on A1 syllable-selective responses are required to distinguishing these two possibilities. Nevertheless, our characterization of distinct types of syllable-selective A1 neurons at the populaton level offers an essential basis for analyzing circuit processing of marmoset vocal sounds.

Neural circuit analysis of complex vocal sounds, including syllables, phrases, and sentences, is beginning to be addressed by advanced technologies that allow recording of population neuronal activity with high spatiotemporal resolution^33–37^. Simultaneous recording of spiking activity in multiple brain regions could further elucidate the spatiotemporal sequence of vocal sound signal processing in unanesthetized animals. In particular, long-term optical recording over large neuronal populations, together with optogenetic manipulation of circuit activity, could help to unravel circuit mechanisms underlying vocal sound processing and experience-dependent circuit plasticity. Develomental and social interaction-dependent changes of marmoset vocal sound production have been observed^38,39^. Whether vocal sound recognition also exhibits plasticity remains unclear. We found no selective response in A1 neurons towards unnatural disyllables when the marmoset was exposed to the latter over periods of minutes. It remains possible that prolonged exposure under appropriate contexts could results in circuit modification that allows marmoset recognition of novel sounds, as suggested by the finding in mice that a sparse set of A1 neurons could become responsive to learned complex sounds^40^.

## Acknowledgements

We thank Yang Dan, Danqian Liu and Xiaoqin Wang for helpful discussion and comments on the manuscript; and Yongheng Fan, Xuebo Li for the care of animals. This work was supported by grants from Chinese Academy of Sciences (No.153D31KYSB20170059, No.XDB32000000, No.QYZDY-SSW-SMCO01), Shanghai Municipal Government (No.2018SHZDZX05, No.18JC1410100, and No.16JC1420201), and Shanghai Post-doctoral Excellence Program (No.2019320).

## Author contributions

H.H.Z., L.P.W. and M.M.P. designed the experiments; H.H.Z. performed the experiments; H.H.Z., J.F.H. and J.R.L. analyzed the data; Z.M.S., N.G. and Y.Q.W. provided technical help; H.H.Z., L.P.W. and M.M.P. wrote the manuscript.

## Data and materials availability

All data necessary to support the paper’s conclusions are present in the main text and Extended Data.

## Methods

### Animals

Animal care and experimental procedures were approved by the Animal Care Committee of Shanghai Institutes for Biological Sciences, Chinese Academy of Sciences. Four adult common marmosets (*Callithrix jacchus*; Marmoset M_a_, M_b_, M_c_ and M_d_; 2 male and 2 females; body weight: 350-430g) provided by the non-human primate facility of the Institute of Neuroscience were used for imaging in this study. Six other marmosets (M_0_ - M_5_) were used for recording call samples. The marmosets were individually housed in a temperature- and humidity-controlled facility (26 - 30⍰°C, 12 h light/dark cycles), and supplied with *ad libitum* water and balanced diet.

### Surgery

Prior to the surgery, the marmoset was first injected with a fentanyl cocktail^6^ intramuscularly (in mg/kg: 0.0005 fentanyl citrate, 0.5 midazolam and 0.05 dexmedetomidine). Afterwards, anesthesia was maintained by 1.5 - 3 % isoflurane with pure oxygen, and the animal was kept in a customized frame in a prone position, with body temperature maintained at ~37⍰°C (monitored with a rectal probe) using a heating blanket. During the surgery, the anesthesia state was confirmed by the absence of pinch-induced paw reflexes. A titanium head-post was attached to the parietal bone by dental cement, performed under sterile conditions. A circular craniotomy (window diameter 8 mm) and durotomy were performed to expose the auditory cortex. The custom-made chronic window consisted of a titanium ring, with coverslip (8 mm in diameter and 0.2 mm in thickness) glued by silicone adhesive (KN-300X, Kanglibang, China), and implanted to the scalp with dental cement seal. The animals were allowed to recover for at least 7 days after surgery. Antibiotics were intramuscularly administered for 3 consecutive days after surgery.

### Cal-520AM Loading

For dye-loading experiments, marmosets were anesthetized by 1.5%-2% isoflurane with pure oxygen through a custom-made mask. The chronic glass window was removed and replaced with a new glass window with a pinhole (~500 μm in diameter), which allowed the cortical access of a dye-loading glass micropipette. For preparing injection solution, Cal-520 AM (40-50 μg, AAT Bioquest) was dissolved with 4 μl Pluronic/DMSO mixture (20% w/v Pluronic F-127 in dimethyl sulfoxide) and was diluted with 35 μl of pipette solution (in mM: 10 HEPES, 2.5 KCl, 150 NaCl, pH 7.4.) and added with 1 μl of Alexa Fluor 594 solution (2 mM) for marking the injected solution. The injection solution was sonicated and filtered with a 0.22 μm filter to remove dye aggregates before loaded into the micropipette (2-4 μm tip opening), and pulse pressure-ejected (3-12 psi, 60-90 pulse at 1 Hz) with a Pico spritzer (Parker Hannifin, USA) to the layer 2/3 of A1 (200-300μm from the surface), as previously described^41^. The surgery and dye ejection usually were performed within 1 hr. After dye ejection, a new glass window without the pinhole was replaced and sealed. The animal was allowed to recover for at least 2 hr before 2-photon calcium imaging was performed when the animal was in the awake state, as indicated by readily licking the milk and eye tracking the experimenter’s action.

### Virus Injection

Injection of GCaMP6f viral vector (for M_b_, M_c_ and M_d_) followed the procedure previously described^11^. In brief, after anesthesia, the chronic window was removed and a glass micropipette (30-μm tip opening) containing 3-4 μl of virus solution was inserted into A1 at a depth of 300-500 μm. The injected viral solution contained the Tetracycline (Tet)-on expression system^11^ consisting of rAAV2/9-hSyn-rtTA (at final titer of 1-3×10^12^ vg/ml) and rAAV2/9-TRE3-GCaMP6f (final titer 2-8×10^12^ vg/ml) as equal-volume mixture. After virus injection (performed within 10 min), a new chronic glass window was installed. The two-photon imaging began ~4 weeks after injection, and GCaMP6f expression was induced by oral Tet administration (doxycycline 0.6 mg/ml in a 5% sucrose solution, 3-5 ml/d) for 3 d prior to the imaging experiment.

### Acoustic stimuli

Marmoset calls were recorded from the marmoset colony at the non-human primate platform of Institute of Neuroscience, Chinese Academy of Sciences, from juvenile or adult marmosets individually in a soundproof chamber (at 48-kHz sampling frequency). Call syllables used in syllable-invariant experiment (Fig. 1) were from marmoset M_1_, M_2_ and M_3_. Four standard test call syllables were sampled from marmoset M_0_ (Phee, Twitter and TrillPhee) and M_5_ (Trill). TrillTwitter was sampled from marmoset M_4_. Call stimuli were presented for 4-5 times with 1-4 s inter-stimulus intervals. For tonotopic mapping with intrinsic optical imaging, pure-tone stimuli (0.5-16 kHz, 4 clicks, 0.2 s duration and 0.5 s interval) were generated using MATLAB (MathWorks) without compression and exposed to the marmoset (48-kHz sampling frequency, 16 bits). The sound delivery system was calibrated using B&K calibrator (2669-L). The sound stimuli were presented by a KEF-ls50 speaker at a distance of 15 cm to the contralateral ear. The sound intensities used in experiments were 70 dB SPL for both pure tones and syllables. The customized sound-insulating chamber used here can attenuate most ambient noise (to a level < 30 dB SPL).

### Spectral-temporal analysis of syllables

To measure the spectral-temporal features of three syllables (Phee, Twitter and TrillPhee) made by conspecific marmosets, we used a customized MATLAB script to analyze 27 calls from 3 marmosets (M_1_, M_2_, M_3_), with 3 calls for each syllable category by each marmoset. We first zero-phase bandpass (3 to 16 kHz) filtered the 27 calls and then measured the following features: duration, amplitude modulation (AM), frequency bandwidth, Wiener entropy, and zero-crossing rate (ZCR). Amplitude modulation was calculated by applying Hilbert transform to call signals to obtain its envelope and further calculating the frequency of the envelope with maximum power. Frequency bandwidth represents the spectrum width of the entire call signal. The bandwidth threshold was set as 1% of the maximum power of the spectrum. Wiener entropy characterizes the width and uniformity of the spectrum and was calculated as the logarithm of the ratio between geometric and arithmetic means of the spectrum^42^. Since ZCRs were different across the call signal, we represented this feature by using the ratio of ZCRs between the first and last 100 ms of the call. Principal component analysis (PCA) was applied to reduce the number of dimensions in the feature, and the first three principal components (PCs) contributed to 98% of the variance.

### Imaging intrinsic optical signals

Marmosets were anesthetized by 1.5 - 2.0 % isoflurane with pure oxygen through a custom-made mask. The head was immobilized with a customized frame. During imaging sessions, the anesthesia was switched to intraperitoneally injection of a fentanyl cocktail (at mg/kg: 0.001 fentanyl citrate, 1 midazolam and 0.1 dexmedetomidine). Signals of reflectance change (intrinsic hemodynamic signals) corresponding to local cortical activity were acquired (Imager 3001, Optical Imaging Inc., Germantown, NY) with 660-nm illumination. Signal-to-noise ratio was enhanced by averaging data from many trials (15–30 trials per stimulus condition). Acoustic stimuli were presented in blocks, with each block containing all frequencies tested or no sound as control. For each condition, imaging began 0.6 s before the sound stimulus onset (for baseline signals). The total imaging time for each frequency was 6 s, during which 30 consecutive frames were collected (at 5 Hz). All stimulus frequencies were displayed in a randomized order.

### In vivo two-photon calcium imaging

Marmosets were progressively trained for about two weeks for habituation in the head-fixed condition in an individually customized frame in the awake state. Fluorescent calcium signals, which are known to correlate with neuronal spiking activity^43, 44^, were monitored from individual A1 neurons with a semi-custom-made LotosScan microscope (LotosScan, Suzhou Institute of Biomedical Engineering and Technology) coupled to a mode-locked Ti:Sa laser (Chameleon VISION-S, Coherent). The excitation wavelength was fixed at 920 nm. Imaging was performed using a 40X, 0.8 NA objective (Nikon). The beam size was large enough to cover the back aperture of the 40X objective. Images were acquired at a frame rate of 40 Hz. In experiment studying the effect of anesthesia, imaging was performed before and one hour after the induction of anesthesia, using the same anesthesia procedure described above.

### Data analysis

Images were analyzed in MATLAB (Mathworks) and ImageJ (National Institutes of Health). For correcting the effect of lateral motion on the imaged data, a rigid-body transformation-based frame-by-frame alignment was applied by using Turboreg plugin (ImageJ software). Neurons were manually identified based on size and shape; astrocytes were identified and excluded based on their distinct morphology^45^. Fluorescence intensity changes with time for each neuron were extracted by averaging pixel intensity values within the cell mask (manually defined) in each frame. Neuropil signal was subtracted by using the method previously reported^46^. After this correction, fluorescence intensity change (ΔF) with time during the stimulus and the post-stimulus period (0.2 s for pure tones, 0.5 s for syllables) were normalized by the pre-stimulus baseline fluorescence (F, over 0.2 s). For each stimulus, the mean ΔF/F was calculated by averaging ΔF/F over the entire stimulus duration for all trials of each stimulus condition. Cells showing significant differences in mean ΔF/F during the baseline *vs*. stimulus-presentation period (*p* < 0.05, ANOVA) were defined as “responsive cells”. Among them, syllable-selective cells were further defined by having responses to some syllables that were significantly higher than those to other syllables (*p* < 0.05, ANOVA) ^47^.

To further confirm the call-selective neurons were exclusively responsive to specific calls rather than statistically biased by the acoustic features compounding the calls, i.e. call frequency band or amplitude which co-varies with the calls, a step-wise GLM (Generalized Linear Model) was conducted. The feature matrix X included eleven predictors which were five acoustic features: duration (s), sound frequency bandwidth (kHz), Wiener entropy, amplitude modulation (Hz) and the ratio over zero from the call onset to offset; and six dummy variables (one-hot) representing the three monkeys’ identity, and the three calls respectively. For each neuron, its activity was taken as the response variable y and a stepwise GLM was run to find out which predictors more significantly contributed to the response variability: starting from a constant model, each time a predictor would be added to the model if it explained the most remaining deviance in the responses. A threshold p<0.05 was taken, and the F-values of the significant predictors per neuron were plotted in Figure 2i (non-significant cases were set to zero) and sorted according to the syllable-selectivity, in descending order. In total, Monkey M_a_ had 550 neurons recorded: acoustic feature cells n = 95, 17.3%, monkey identity cells n = 30, 5.5%, call cells n = 38, 6.9% (among them 27 cells were purely call selective, namely non-significant to identity and acoustic features); Monkey M_b_ had 468 neurons recorded: acoustic feature cells n = 93, 19.9%, monkey identity cells n = 28, 6%, call cells n = 43, 9.2% (among them 22 cells were purely call selective, namely non-significant to identity and acoustic features). To visualize the neural representation of each sub-group of neurons on low dimensions, MDS (Multi-Dimensional Scaling) was used. It calculated the distances (Euclidean distance) between the sample points on the high dimensional space, and projected the samples on a 2- or 3- low dimensional space.

**Syllable Selectivity Index (SSI)** was calculated based on the equation SSI = (R_pref_−R_nonpref_)/(R_pref_+R_nonpref_), where R_pref_ and R_nonpref_ represent the mean ΔF/F evoked by the preferred syllable and all other syllables, respectively. The **Modulation Index (MI)** was calculated based on peak ΔF/F values by the equations below (Phee cell as an example): MI = (R_TwP_−R_P_)/(R_TwP_+R_P_), where R_TwP_ and R_p_ represent the average peak ΔF/F responses evoked by TwitterPhee and Phee, respectively. To analyze the nearest-neighbor distances for all syllable-selective cells regardless of syllable selectivity, a bootstrap sampling method was used. For each trial of sampling, the same number of neurons as in each syllable-selective ensemble was randomly selected from the syllable-selective populations in each imaging field. We used 500 sampling trials and obtained an averaged cumulative percentage plot of nearest-neighbor distances.

## Code and data availability

Custom MATLAB scripts used to analyze and plot all data, and original data collected in this study are available from the corresponding authors upon request.

---

**Extended Data Figure 1.**
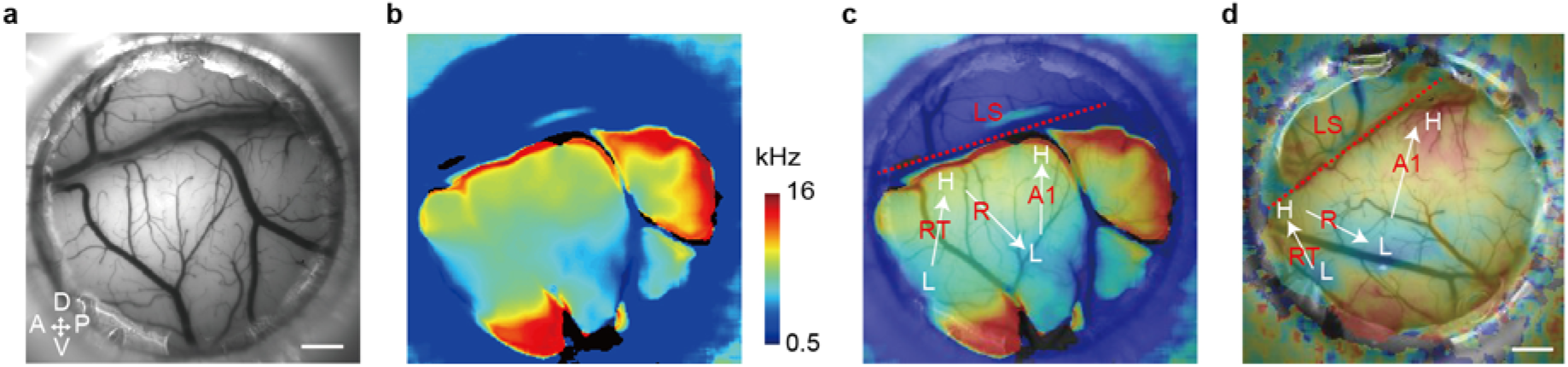
Tonotopic map of the auditory cortex obtained by imaging intrinsic optical signals. **a**, Blood vessel map within the imaging window (M_a_). **b**, A tonotopic map revealed by intrinsic optical signals in response to a sequence of 21 discrete pure tone stimuli in the range of 0.5-16 kHz. The frequency preference of different imaged regions was color-coded by the scale bar. **c**, Image obtained by merging images in **a** and **b**. **d**, Same as **c** for another marmoset (M_d_). LS, lateral sulcus; A1, primary auditory cortex; R, rostral field; RT, rostro-temporal field. Bars: 1 mm.

**Extended Data Figure 2.**
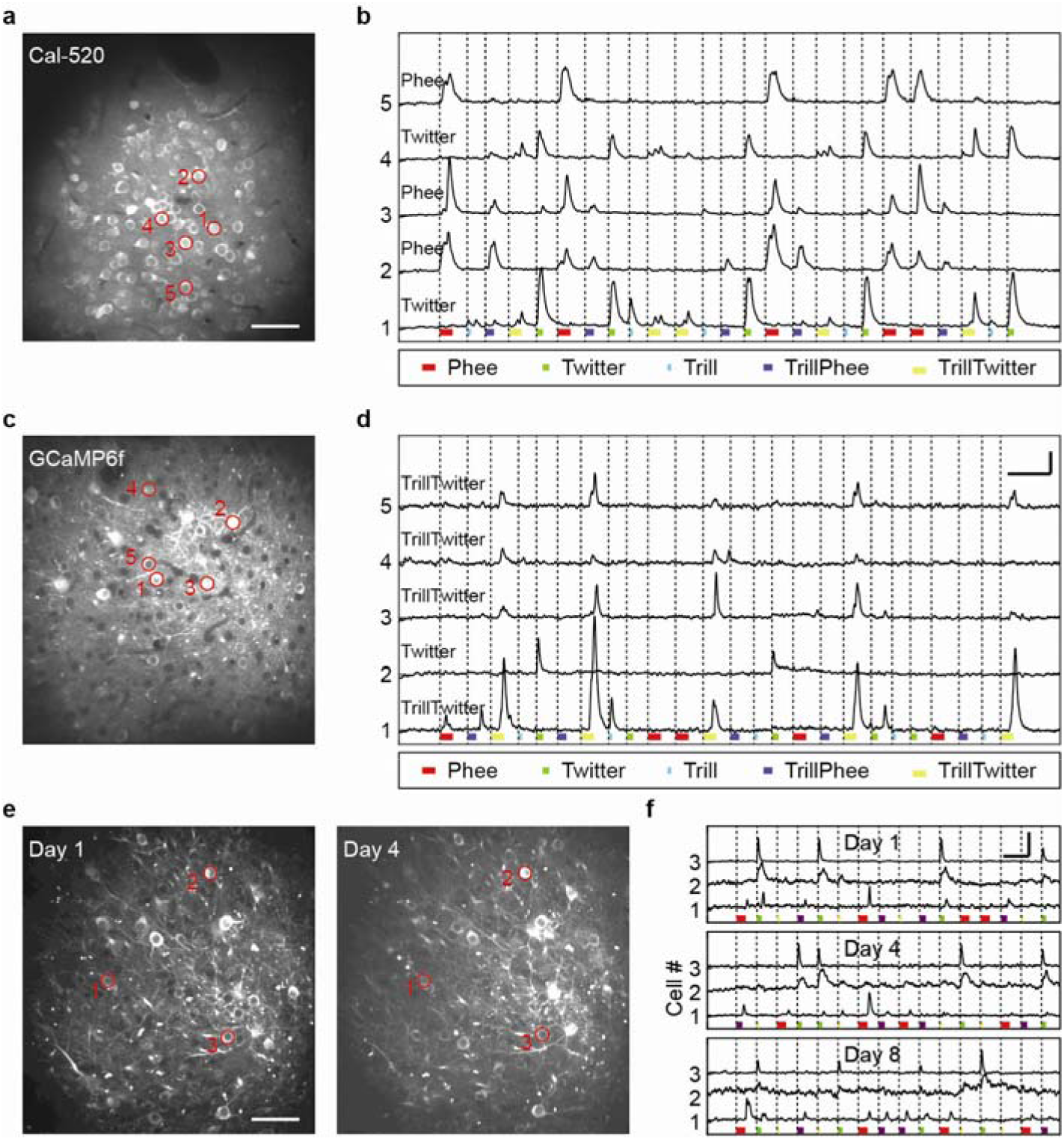
Two-photon imaging of neuronal activity in awake marmoset A1. **a**, Fluorescence image of an A1 area of marmoset M_a_, loaded with Cal-520AM. Bar: 50 μm. **b**, Relative changes in fluorescence (ΔF/F) in 5 example cells (marked by circles in **a**) in response to 5 different call syllables in a random sequence. Stimulus duration marked by the bar below, syllable types coded in colors. The apparent syllable selectivity was assigned to each cell (marked on the trace). **c, d**, Similar to **a** and **b**, except that the A1 area was injected with Tet-dependent AAV expressing GCaMP6f. Data from marmoset M_a_. Bars: 5 s and 100% ΔF/F. **e**, Fluorescence image of an A1 area of a GCaMP6f-expressing marmoset (M_d_) on day 1 and day 4 of the experiment, which began 35 days after AAV injection, and 3 days after Tet application. Bar: 50 μm. **f**, Response profiles of 3 example neurons, marked by red circles in e and recorded at day 1, 4 and 8, showing the same apparent syllable selectivity. Bars: 5 s and normalized ΔF/F.

**Extended Data Figure 3.**
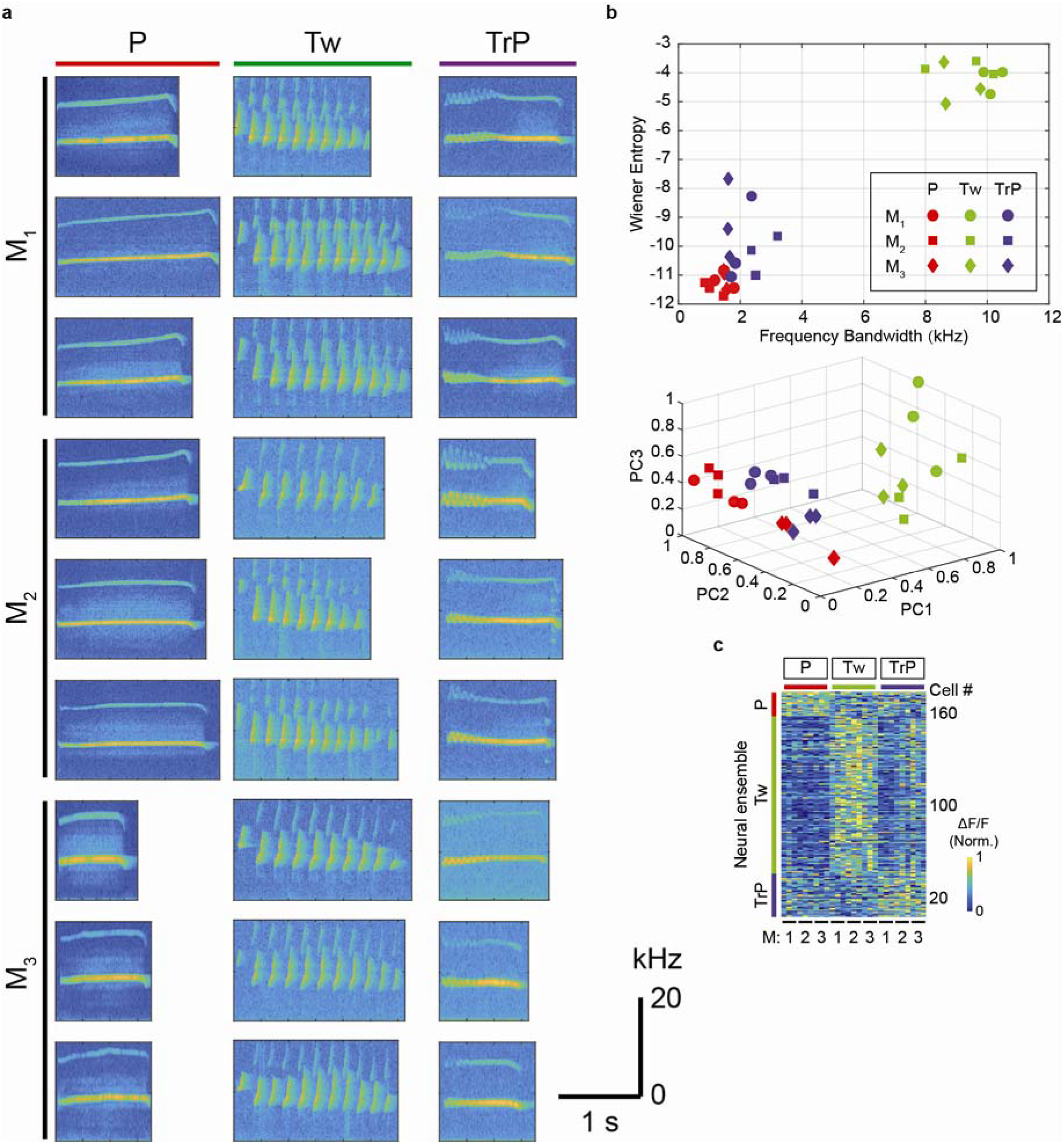
Spectrograms of all 27 test call syllables from 3 marmosets (M_1_, M_2_, M_3_). **a**, Spectrograms of 27 syllable stimuli. **b**, Bandwidth Wiener-entropy analysis and principal component analysis (PCA) of 27 syllable stimuli shown in **a**. Shown in the bottom is first three principal components (PC). **c**, Heat map for all syllable responsive neurons in marmoset M_b_ that was exposed to three syllable categories as in a. Each horizontal line depicts average amplitude of ΔF/F (from 5 trials), with 3 representative calls from each marmoset for each syllable. The cells were sorted into three syllable ensembles, based on the syllable category that evoked the maximal mean ΔF/F amplitude. The amplitude is coded in color by the scale shown on the right.

**Extended Data Figure 4.**
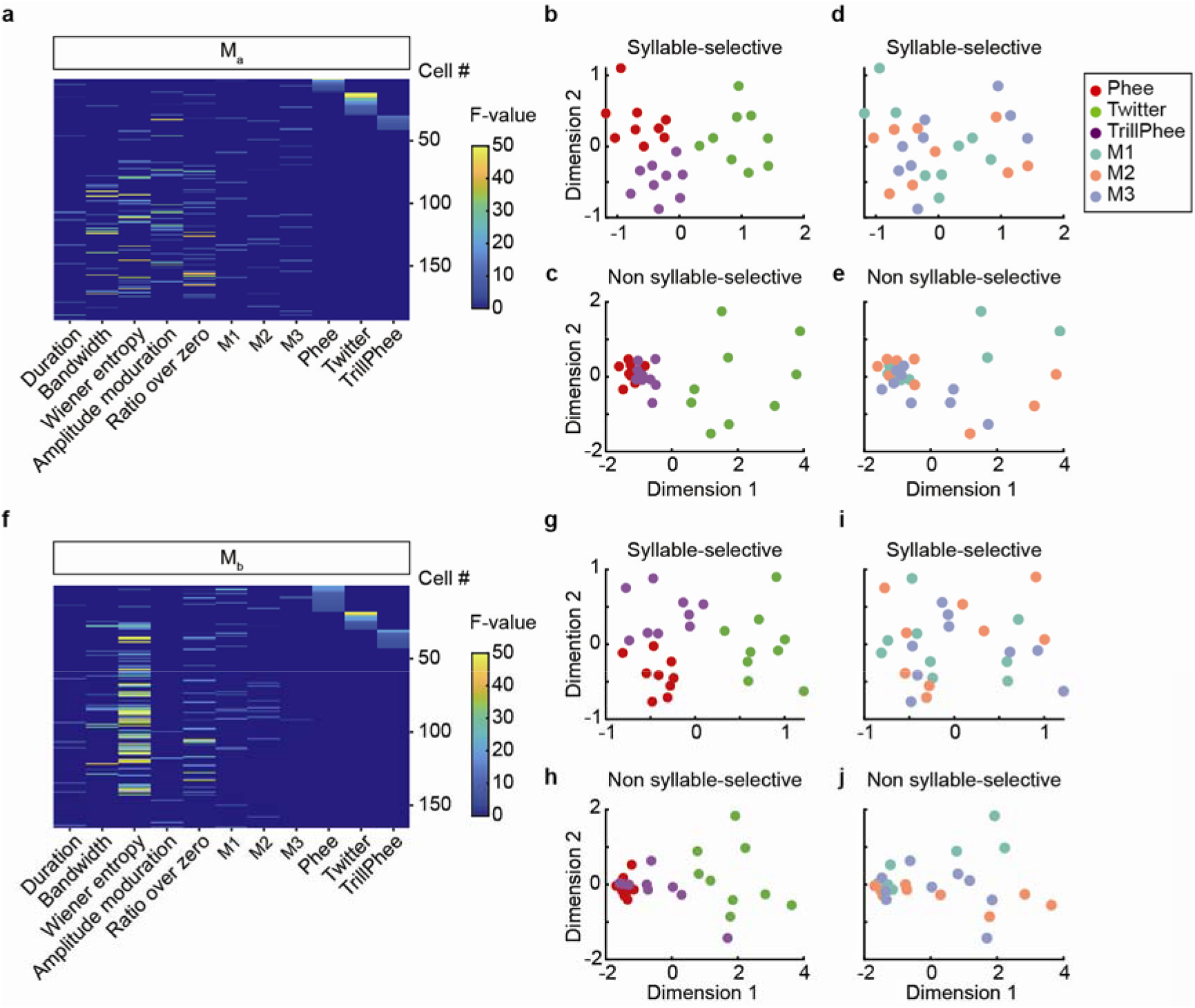
a, Quantification of the relative selectivity of syllable-selective cells to specific acoustic, animal identity, and syllable variables in the stimulus, using a generalized linear model (see Methods). For each variable, the column depicts the significance level for response modulation by that variable of all neurons from Ma recorded in experiments shown in Fig. 1. The neurons were sorted according to their syllable selectivity. **b**, Multi-dimensional scaling plot of neuronal representation of the 27 syllables (shown only for two dimensions) by the subsets of the neurons depicted in **a**, based on their selectivity: neurons that responded exclusively to syllables and invariant to other variables (n = 27/193); syllable-selective neurons that also responded to non-syllable variables (**c**, n = 90/193). **d** & **e**, Same neural representations using the label of animal caller disrupted such neural clustering. Each dot denotes one syllable. **f-j**, Same depiction as **a-e**, except that the data were from Mb (n = 22/165, g; n = 80/165, h). Note distinct clusters for three syllables in neurons that responded exclusively to syllable variable in both marmosets.

**Extended Data Figure 5.**
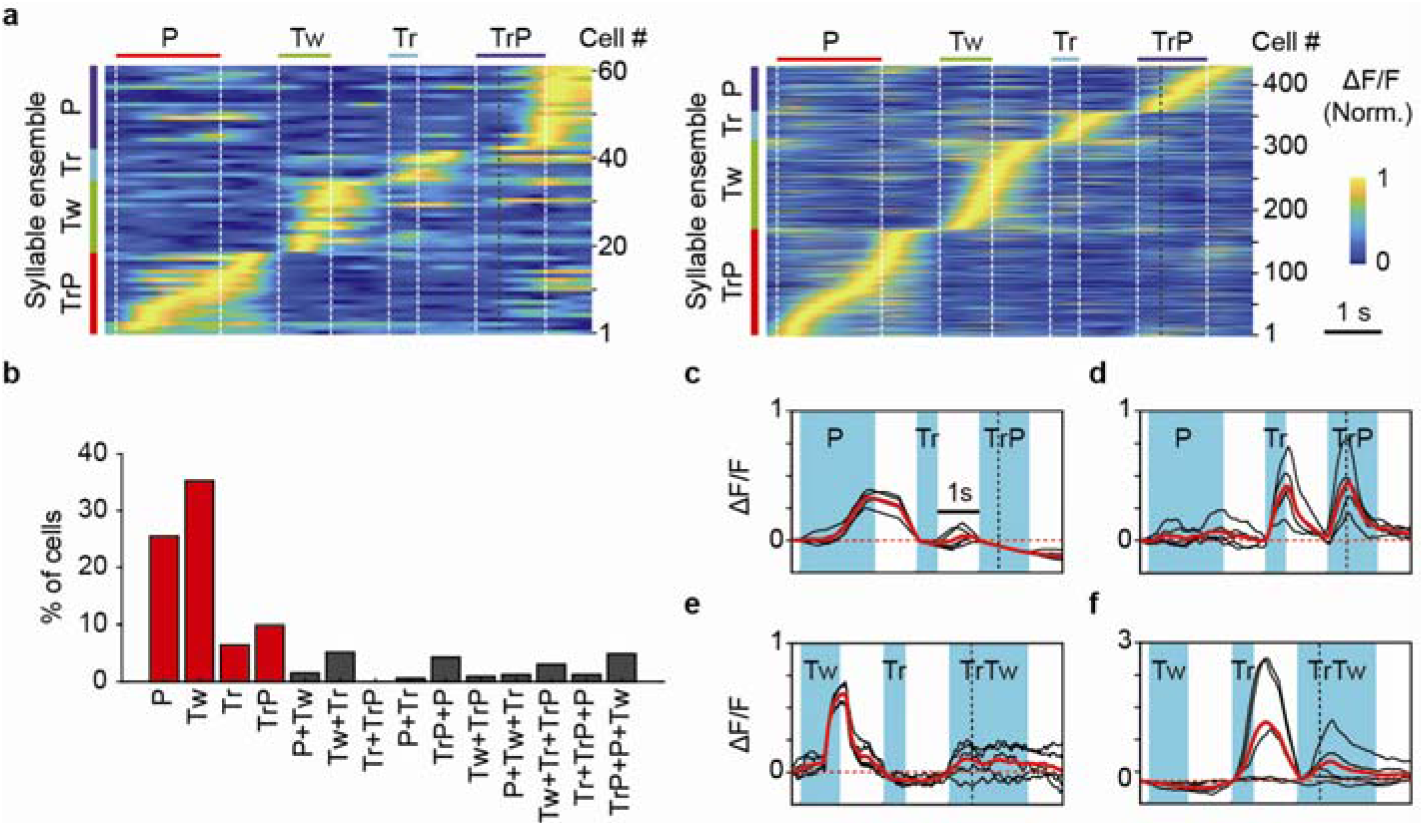
Summary of syllable-selective neuronal ensembles from two other marmosets recorded (M_a_, M_d_). **a**, Activities of all syllable-selective neurons recorded from marmoset M_d_ (left) and M_a_ (right) labeled with Cal-520AM, shown by the heat map in the same manner as in Fig. 1c. **b**, Percentages of cells showing selectivity to single syllable and to different sets of multiple syllables, among all syllable-selective cells. **c** and **d**, Fluorescence changes (ΔF/F; black, individual trials; red, average) evoked by disyllable TrillPhee (TrP) and its component monosyllable Phee (P) and Trill (Tr), in two neurons recorded from marmoset M_c_. Red dash lines indicate zero ΔF/F. Note that this cell (**c**) exhibits negative ΔF/F to TrillPhee. **e** and **f**, Similar to **c** and **d**, but for responses evoked by TrillTwitter (TrTw), Trill (Tr) and Twitter (Tw). Neurons recorded from marmoset M_c_.

**Extended Data Figure 6.**
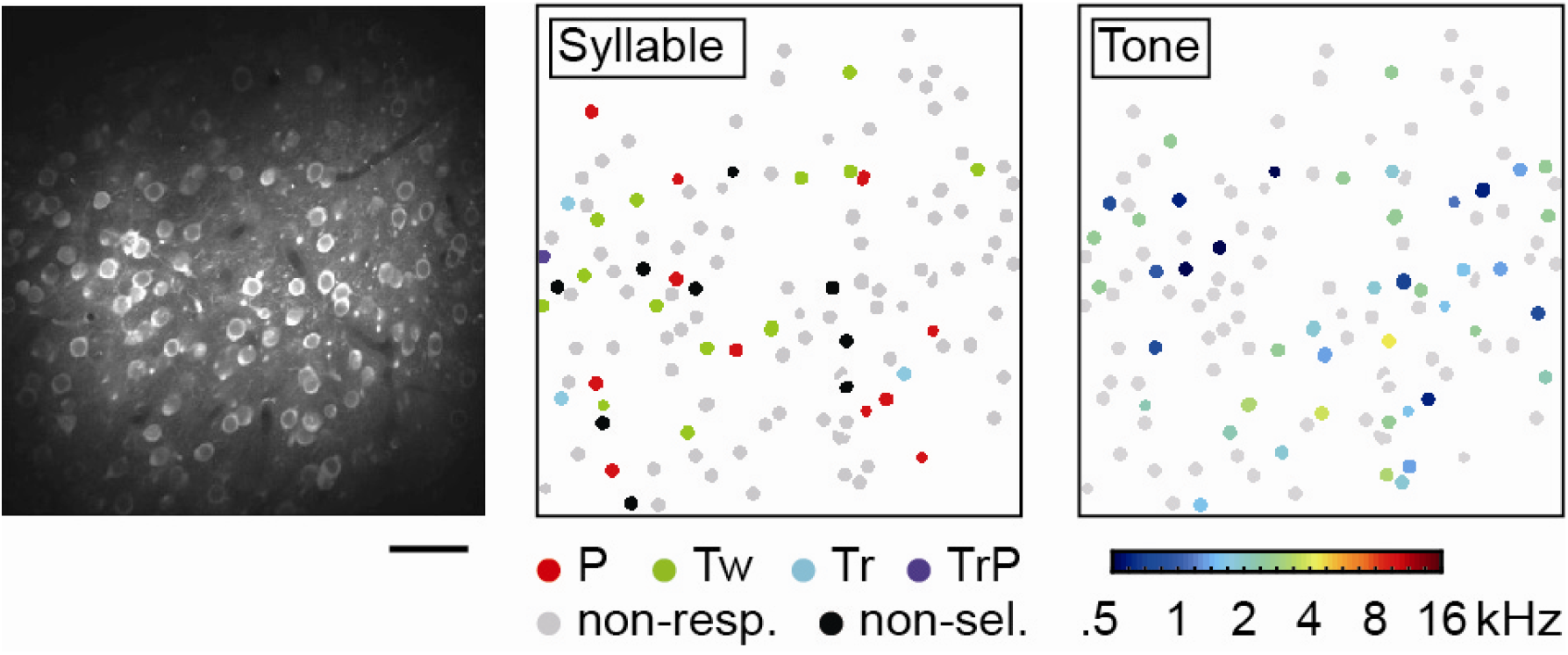
Responses to syllables and pure tones of A1 neuronal populations in 2 kHz area. Left, an image of Cal-520AM fluorescence at a recorded region of marmoset M_c_. Bar: 50 μm. Middle, spatial distribution of all syllable-selective cells in the imaging field, with response properties (“non-resp.”, not responding to syllables; “non-sel.”, non-selective responses to syllables) coded in colors. Right, tone-selective cells within the same imaging field, with tone frequency coded in color scale shown below.

**Extended Data Figure 7.**
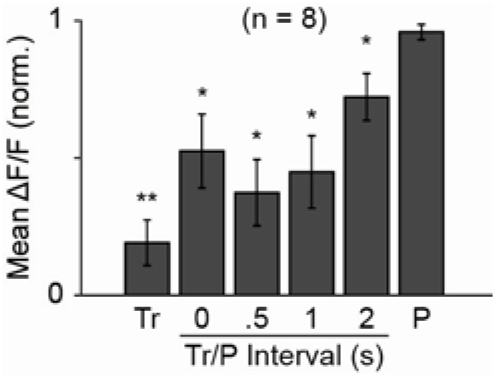
Suppression of Phee-evoked responses of Phee-selective monosyllable neurons by a preceding Trill isolated from a natural TrillPhee call. The data represent mean peak ΔF/F values evoked in Phee-selective neurons, observed when the Trill/Phee interval was set at 0, 0.5, 1.0 and 2.0 s, together with responses evoked by Trill alone in these Phee-selective neurons (error bar, SEM; n = 8 cells). The mean amplitudes for Trill- or Trill/Phee-evoked responses were significantly smaller than those evoked by Phee alone (paired t test; **, *P* < 0.001; *, *P* < 0.05).

