## Supplementary figures and images for "Distinct natural syllable-selective neuronal ensembles in the primary auditory cortex of awake marmosets"

### Extended Data Figure 1

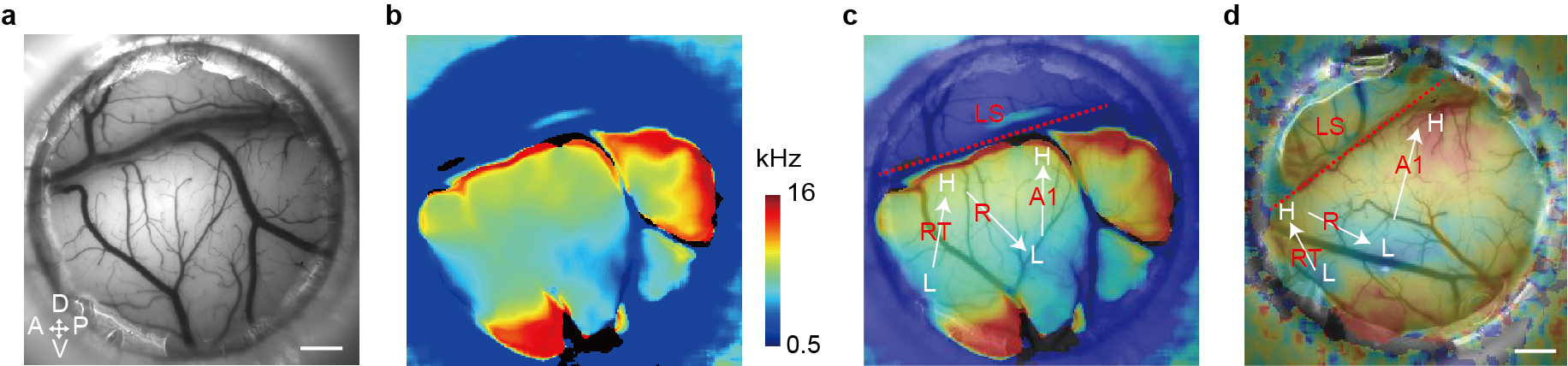

### Extended Data Figure 2

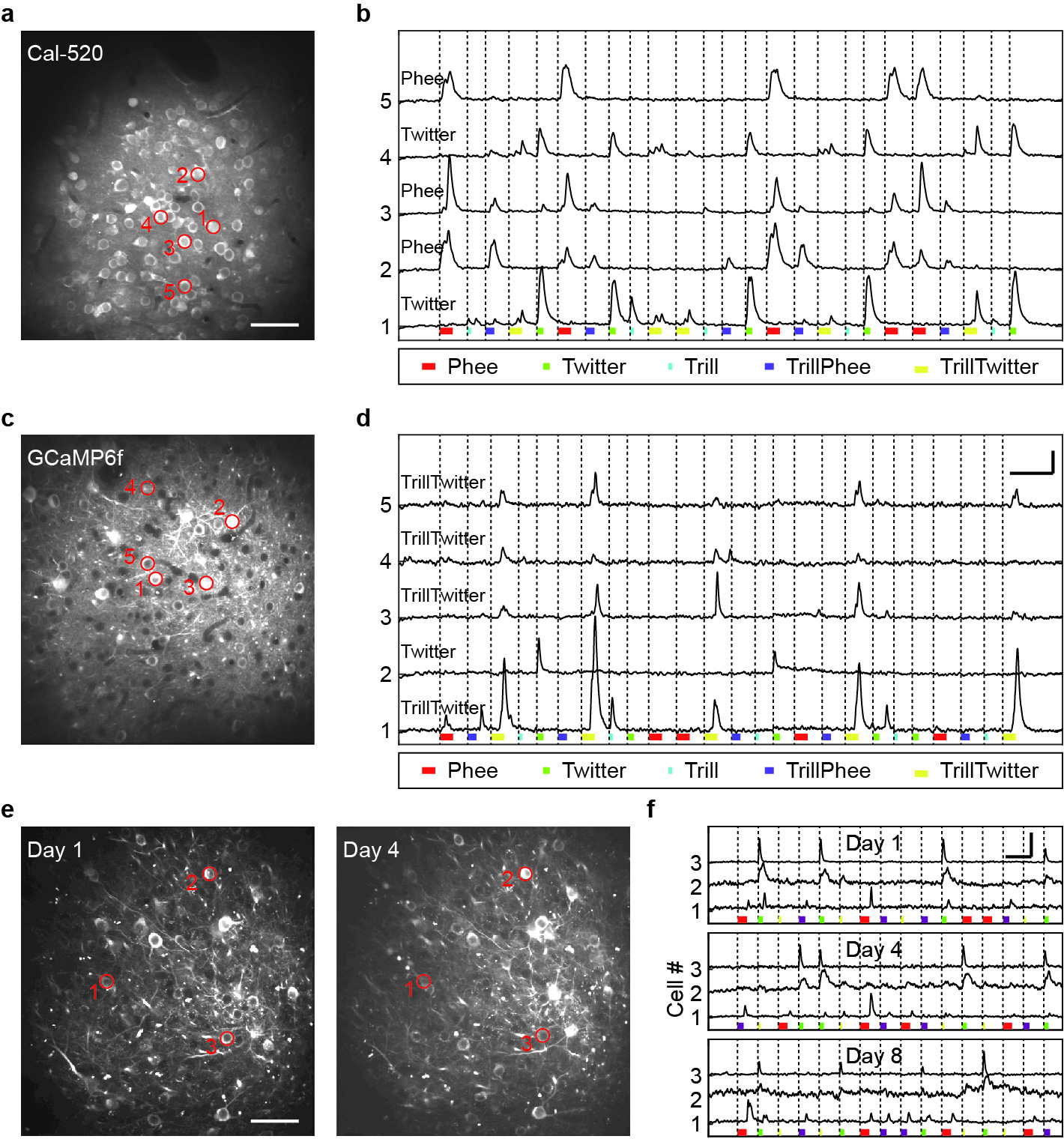

### Extended Data Figure 3

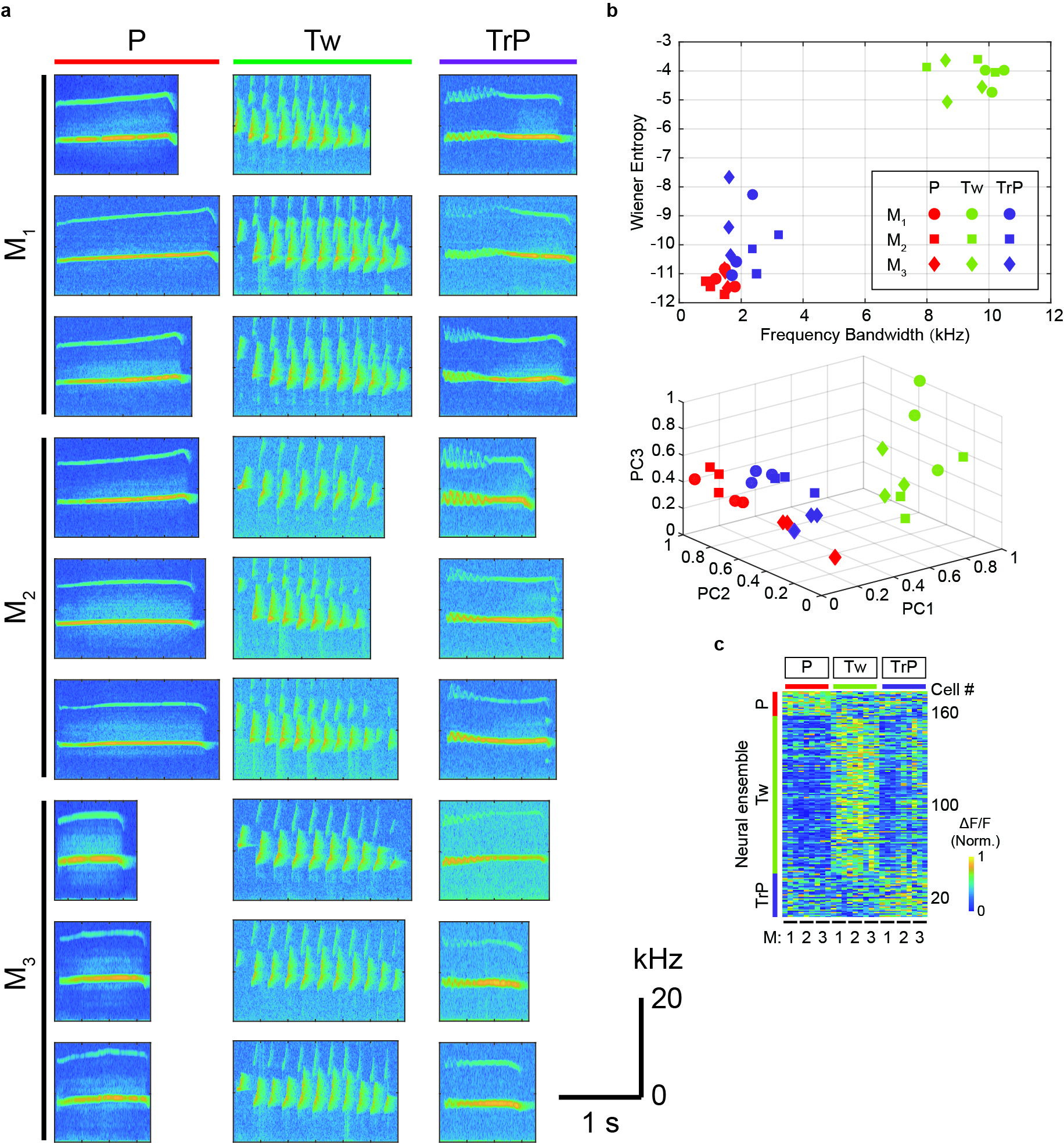

### Extended Data Figure 4

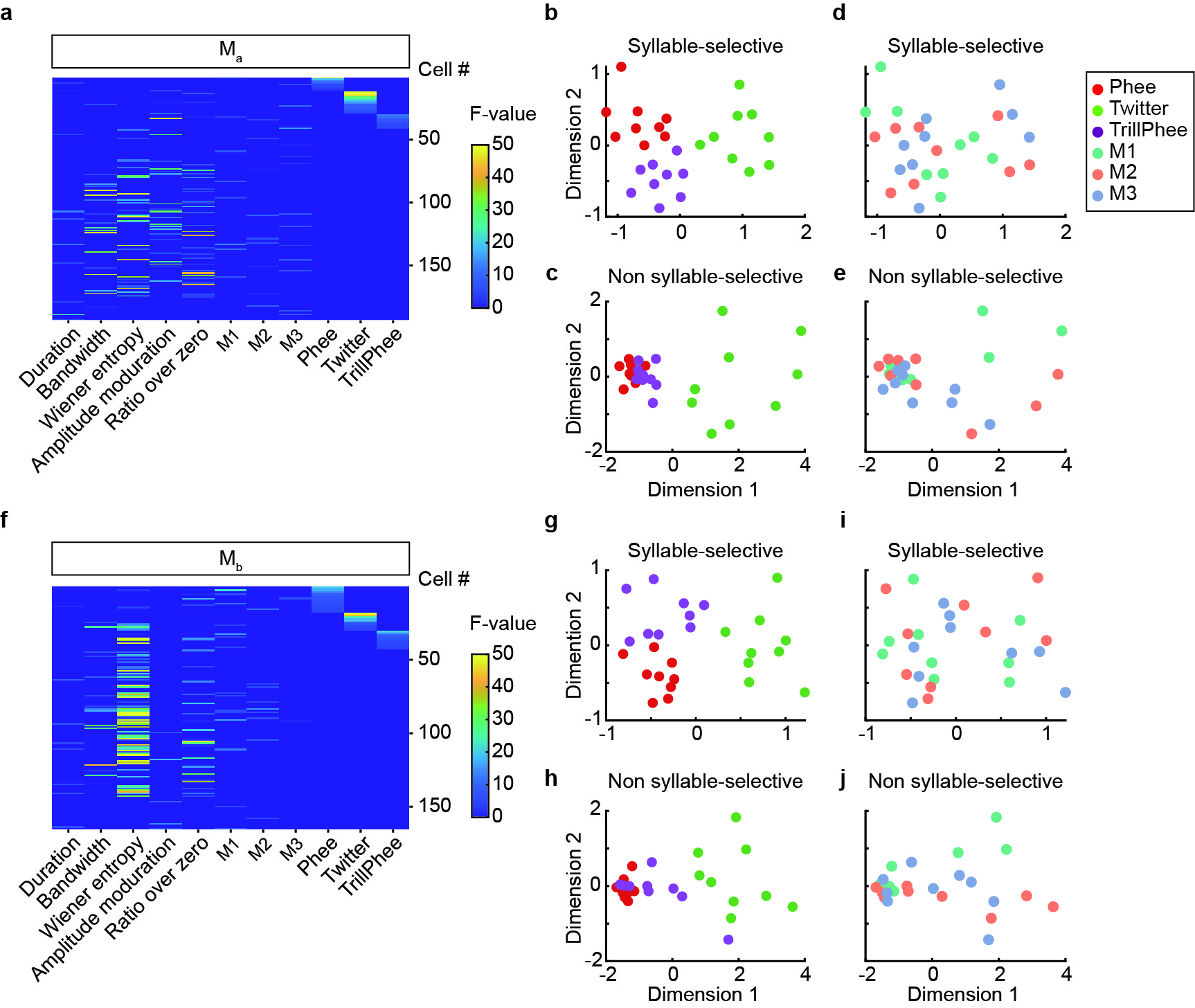

### Extended Data Figure 5

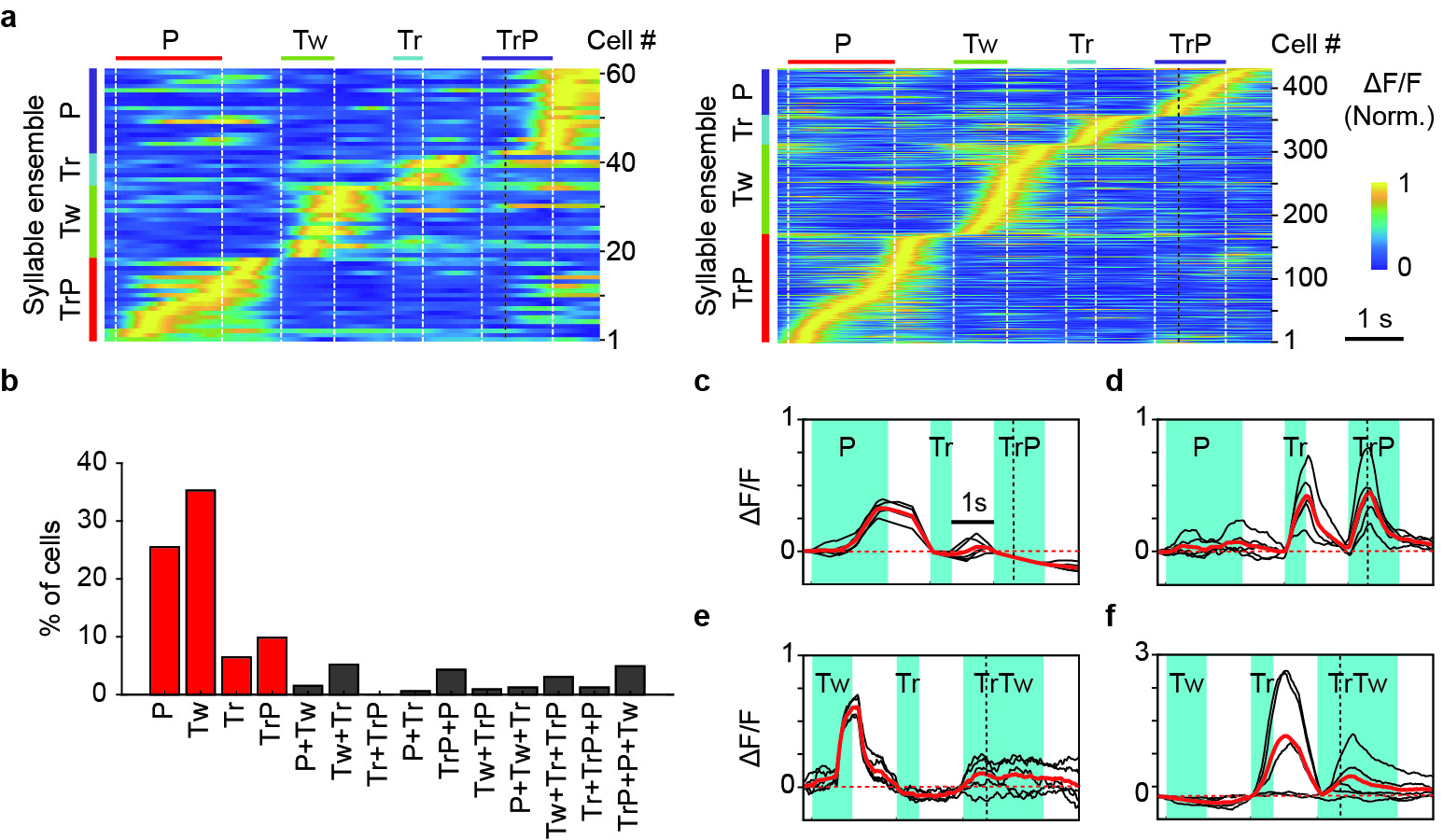

### Extended Data Figure 6

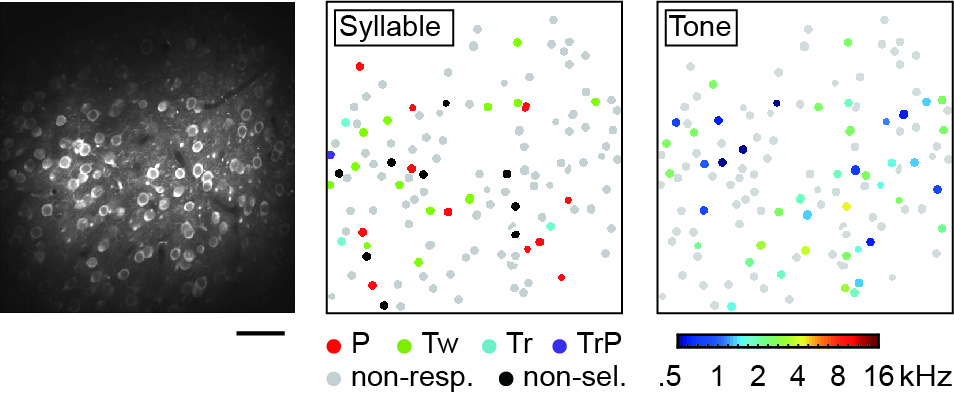

### Extended Data Figure 7

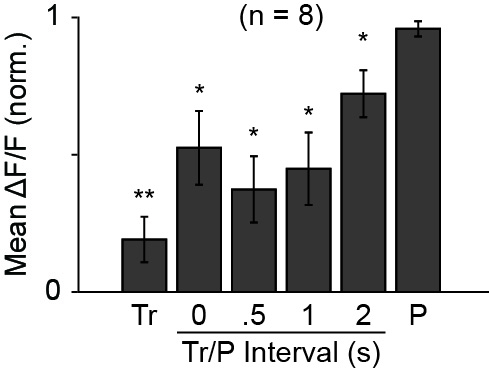
